# Mutational signatures of complex genomic rearrangements in human cancer

**DOI:** 10.1101/2021.05.16.444385

**Authors:** Lisui Bao, Xiaoming Zhong, Yang Yang, Lixing Yang

## Abstract

Complex genomic rearrangements (CGRs) are common in cancer and are known to form via two aberrant cellular structures—micronuclei and chromatin bridge. However, which mechanism is more relevant to CGR formation in cancer cells and whether there are other undiscovered mechanisms remain open questions. Here, we analyze 2,014 CGRs from 2,428 whole-genome sequenced tumors and deconvolute six CGR signatures based on the topology of CGRs. Through rigorous benchmarking, we show that our CGR signatures are highly accurate and biologically meaningful. Three signatures can be attributed to known biological processes—micronuclei- and chromatin-bridge-induced chromothripsis and extrachromosomal DNA. More than half of the CGRs belong to the remaining three newly discovered signatures. A unique signature (we named “hourglass chromothripsis”) with highly localized breakpoints and small amount of DNA loss is abundant in prostate cancer. Through genetic association analysis, we find *SPOP* as a candidate gene causing hourglass chromothripsis and playing important role in maintaining genome integrity. Our study offers valuable insights into the formation of CGRs.

## Introduction

Genome instability is a hallmark of cancer^1,2^, and somatic genome rearrangements are abundant in human cancers^3,4^. Genomic rearrangements, also known as structural variations (SVs), include simple forms such as deletions, duplications, inversions and translocations, as well as complex forms^3,5,6^. A peculiar type of extremely complex genomic rearrangements (CGRs) called chromothripsis refers to a single catastrophic event resulting in numerous somatic genome rearrangements, and has been found in many tumor types^7,8^. In addition, other forms of CGRs have been described, such as chromoanasythesis^9^, chromoanagenesis^10^ and chromoplexy^11^.

Understanding the molecular mechanisms leading to the formation of somatic genome rearrangements in cancer is of clinical significance in disease screening^12^ and treatment^13^. Different mutational processes operate in different tissues and leave distinct footprints (genetic alterations) in the DNA which can be used to mathematically decompose the mutational signatures corresponding to individual mutational processes. Mutational signatures have previously been deconvoluted for somatic alterations, including single nucleotide variants (SNVs)^14^, copy number variations (CNVs)^15^ and SVs^16,17^. Although CGRs are abundant in cancer^8^, their forming mechanisms are still largely unknown. We hypothesize that multiple molecular mechanisms can lead to CGR formation and the mechanisms can be inferred based on the patterns of CGRs. Recently, several studies have successfully recapitulated chromothripsis *in vitro* and have revealed at least two CGR forming mechanisms with distinct patterns. The first mechanism is through the formation of micronuclei due to chromosomal segregation errors^18^. In this mechanism, chromosomes trapped in micronuclei shatter into many pieces and randomly rejoin^19^. The rearrangement breakpoints are evenly distributed across the chromosomes and the DNA segments have two or three copy-number states^18^. The second mechanism is through chromatin bridge formed by dicentric chromosomes^20,21^. The bridge breaks due to mechanical force and results in chromothripsis^22^ with highly localized breakpionts^20,22^. Chromatin-bridge-induced chromothripsis also displays two or three copy-number states. Before being resolved as chromothripsis, the dicentric chromosome may undergo breakage-fusion-bridge (BFB) cycles during which DNA fragments can be duplicated and lost. Therefore, the copy-number states in the BFB-cycle-induced chromothripsis can be more than three. In BFB-cycles/chromatin-bridge-induced chromothripsis, the chromosomes involved often lose telomeres, and foldback inversions are often present. Moreover, micronuclei and chromatin bridge can both lead to the formation of extrachromosomal DNA^18,23^ (ecDNA), a circular form of DNA molecules known as double minutes (DMs), which is common in some tumor types^24^. In ecDNA, small fragments of DNA from different regions of the genome are often connected and highly amplified. However, the extent to which of these mechanisms contribute to CGR formation in disease tissues from patients, and whether there are additional mechanisms of CGR remain unclear.

Methods such as non-negative matrix factorization have been developed to decompose the mutational signatures^14^. However, such matrix-factorization-based strategy cannot be used to decompose signatures of CGRs, because a large number of variants are required in the tumor genomes. Although numerous rearrangements are present in CGRs, they are formed as a one-time event, and each tumor only carries one or two of such events. Therefore, new approaches are needed to study CGR signatures. Several studies have used graphs to classify CGRs^25,26^, but had limited abilities to incorporate the breakpoint distribution and telomere loss aspects of CGRs. Here, we describe a robust computational algorithm called “Starfish” to deconvolute CGR signatures in human cancer based on the topology of CGR events.

## Results

### CGR signatures based on event topology

The definition of chromothripsis has been imprecise and evolving^7,27^. A previous review^27^ proposed a set of criteria to define chromothripsis: clustering of breakpoints, randomness of DNA fragments, oscillating copy-number states, involvement of a single haplotype and ability to walk through the derived chromosome, etc. In our recent study^8^, we followed the above criteria to detect chromothripsis using ShatterSeek in 2,428 whole-genome sequenced tumors from Pan-cancer Analysis of Whole Genomes (PCAWG). However, in some studies, the term chromothripsis is used loosely without the requirement of oscillating copy-number states^23,28^. Here, we define “CGRs” broadly as complex events formed via one-time events rather than accumulation of multiple individual events over time. In order to detect CGRs more broadly, we removed the requirement of oscillating copy-number states in ShatterSeek (see details in Online Methods) and re-analyzed the same 2,428 samples^8^. A total of 2,014 CGRs were detected in 1,289 (53.1%) samples (Supplementary Table S1). The newly detected 1,226 CGRs demonstrated interleaved SVs (Supplementary Figure S1) which suggested that they likely formed through one-time catastrophic events. Out of 285,791 somatic SVs detected in 2,428 tumors, 106,759 (37%) are involved in CGRs. Therefore, CGRs are a major source of genomic instability.

We hypothesize that the CGR topology—pattern of copy-number states and distribution of breakpoint junctions—can be used to infer their mechanisms of formation. To this end, we developed a computational algorithm “Starfish” to deconvolute CGR signatures based on the event topology (see details in Online Methods). Briefly, we first selected 12 features to comprehensively depict the topology of each CGR event (Figure 1a). These features include breakpoint dispersion score, copy loss percentage, copy loss density, copy gain percentage, copy gain density, number of copy states, median copy number change, maximum copy number, highest telomere loss percentage, ratio of telomere loss and CGR loss sizes, median breakpoint microhomology and median breakpoint insertion size. Features related to the magnitude of CGR events (i.e. number of chromosomes involved, number of rearrangements and size of CGR regions) were not used. After removing highly correlated features (Supplementary Figure S2a) and features with small variances (Supplementary Figure S2b), there were five features remaining: CGR breakpoint dispersion score (measuring the randomness of breakpoint distribution on chromosomes), copy loss percentage, copy gain percentage, telomere loss percentage and maximum copy number. We performed unsupervised consensus clustering for all 2,014 CGRs (Figure 1a) using the five features and discovered six clusters (Figure 1b, Supplementary Figure S2c). Clusters constructed by different clustering approaches were very similar (Supplementary Figure S2d). We use six clusters produced by the partition around medoids (PAM) algorithm for the remainder of this document and refer to the clusters as CGR signatures (Supplementary Table S1).

Signature 1 features highest maximum copy number with moderate percentages of copy gain (Figure 1b). This pattern resembles ecDNA where small DNA fragments are highly amplified (Figure 1c). Homogeneous staining region (HSR) is another form of genomic amplification in cancer and the amplified DNA fragments reside on linear chromosomes^29^. HSRs can excise from the host chromosomes and circularize into DMs and DMs can re-integrate into linear chromosomes^23,30^. The Signature 1 does not differentiate between circular and linear forms, and therefore, it captures both DMs and HSRs. Signature 2 has the highest amount of telomere loss (Figure 1b) and reflects chromothripsis formed through chromatin bridge and possibly involving BFB cycles (Figure 1c). Signatures 3 and 5 present the largest amount of genomic copy loss and copy gain respectively (Figure 1b and 1c) and neither matches any known mechanisms. Signature 4 highlights the lowest breakpoint dispersion scores (breakpoints evenly distributed) with modest copy gains and losses (Figure 1b) which coincides with micronuclei-induced chromothripsis (Figure 1c). In sharp contrast, Signature 6 manifests the highest breakpoint dispersion scores (breakpoints unevenly distributed) with a very small amount of copy loss and no copy gain (Figure 1b). This pattern fits the definition of chromothripsis^27^ with distinct patterns, so we named this signature “hourglass chromothripsis” (Figure 1c) due to the shape of its copy-number profile (small fractions of content leaking to the lower level). Among the six signatures, Signature 3 is the least common one with 240 events and Signature 6 is the most abundant one with 431 events (Figure 1b). In total, 971 (48%) CGRs can be related to known mechanisms (Signatures 2 and 4) and known cellular structures (Signature 1). Interestingly, 1,043 (52%) CGRs belong to the three signatures (Signatures 3, 5 and 6) that cannot be attributed to any known mechanisms which highlights the benefit of our signature decomposition strategy.

**Figure 1.**
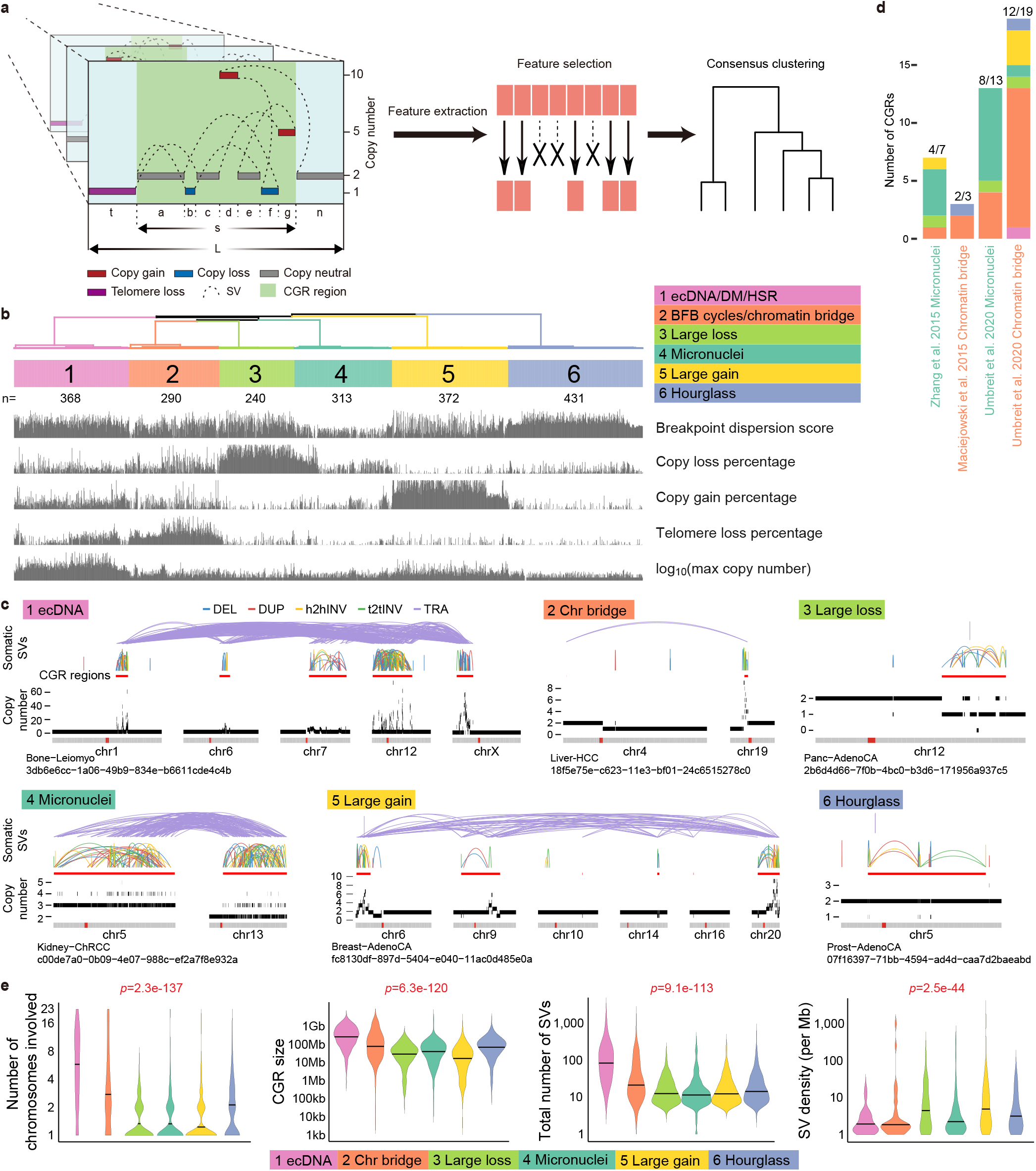
Six CGR signatures in PCAWG cohort. **a**, Starfish workflow. CGR regions are identified in 2,428 PCAWG samples. Twelve features are selected to comprehensively describe the topology of CGR events. Highly correlated features and features with low variability are removed. The remaining features are used to perform unsupervised clustering. Letters “a” to “g”, “n”, “t”, “s” and “L” denote the lengths of DNA segments. **b**, Six CGR signatures (clusters) based on five features. **c**, Examples of CGRs from six signatures. In each example, chromosomes involved in CGRs are displayed side-by-side. The arcs in top panels are five types of somatic SVs (deletions, duplications, head-to-head inversion, tail-to-tail inversions and translocations). CGR regions are marked by red bars below the SV arcs. The bottom panels are copy number profiles. **d**, Benchmarking CGR signatures using experimentally induced events. The texts of the x axis are colored based on experimental CGR forming mechanisms. Starfish predicted CGR signatures are shown as bars **e**, Differences in magnitude of six CGR signatures. Horizontal lines in violin plots are median values. *P* values are calculated by Kruskal–Wallis tests.

### Benchmarking CGR Signatures

To benchmark our algorithm, we utilized three studies that have induced chromothripsis events using experimental approaches^18,20,22^. In order to easily classify additional CGR events, we trained a neural network classifier (Supplementary Figure S3a and S3b), namely Starfish classifier, using the five features and CGR signature labels from the PCAWG samples (Figure 1b). We then predicted CGRs and their signatures in the three benchmarking datasets using the modified ShatterSeek and Starfish classifier. In 62% of the cases (26 out of 42, Supplementary Table S2), the predicted signatures matched experimentally induced CGR forming mechanisms (Figure 1d) which suggested that the CGR signatures deconvoluted by Starfish are biologically meaningful. In the study by Umbreit et al., micronuclei were formed via broken chromatin bridge^22^ and produced chromothripsis through DNA shattering. Consistent with this, most CGRs (8 out of 13) in this category displayed patterns of micronuclei-induced chromothripsis (Figure 1d).

As an indirect validation of Signature 1, we compared Signature 1 CGRs to ecDNA (circular events) predicted by AmpliconArchitect^25^. Since 85% of the AmpliconArchitect-predicted circular events in cell lines were confirmed to be DMs by florescent in situ hybridization (FISH)^25^, we used them as ground truth. In 289 AmpliconArchitect-predicted circular events from 849 tumors shared between this study and the AmpliconArchitect study, most were relatively simple (Supplementary Figure S3c) as ecDNA/DM can form with only one DNA fragment ligated head-to-tail^3,23^. Out of the 100 complex circular events, 64 (64%) were classified as Signature 1 by Starfish. There were also 309 CGRs in Signature 1 not classified as circular events by AmpliconArchitect which could be the linear form of HSRs. To validate this, we compared them to tyfonas events predicted by JaBbA^26^ which were experimentally validated as HSRs. In 887 tumors shared by this study and JaBbA study^26^, 38 out of 46 (83%) JaBbA-predicted tyfonas events were classified as Signature 1 by Starfish (Supplementary Figure S3d). These results combined suggested that the Signature 1 can truthfully capture both DMs and HSRs.

In summary, our CGR signatures deconvoluted based on event topology are highly accurate and biologically relevant.

We then investigated the differences in event magnitude among six CGR signatures. Signatures 1 and 2 have more chromosomes and SVs involved and affect larger genomic regions compared to the other signatures (Figure 1e). The number of foldback inversions has been used to classify BFB-cycle events^25,26^. However, we found that a hard cutoff with the number of foldback inversions cannot effectively separate Signature 2 from others (Supplementary Figure S3e).

### CGR events as major drivers of cancer

The frequencies of CGRs significantly vary across tumor types (Figure 2a and Supplementary Figure S4a). CGRs are most abundant in glioblastoma, osteosarcoma and esophageal cancer, while pilocytic astrocytoma and chronic lymphocytic leukemia barely have any. Signature 1 is frequently observed in many tumor types including soft tissue sarcoma, esophageal cancer, glioblastoma and lung squamous cell carcinoma (Figure 2a and Supplementary Figure S4b), which is consistent with previous studies^24,25^. In contrast, Signature 2 is most abundant in clear cell renal cell carcinoma (Figure 2a and Supplementary Figure S4b) in which chromosomes 3 and 5 are known to be prone to chromothripsis^31^. This signature is also abundant in osteosarcoma, melanoma, breast cancer, and ovarian cancer (Figure 2a). The enrichment of the Signatures 1 and 2 being consistent with previous studies again demonstrated the accuracy of our signature deconvolution. Strikingly, Signature 6 is found in almost all tumor types (Figure 2a) and is particularly common in prostate cancer (107/187 [57%] in PCAWG cohort). We also observed biases of CGR occurrences among tumor subtypes. For example, Signature 5 is enriched in basal breast cancers (Supplementary Figure S4c).

**Figure 2.**
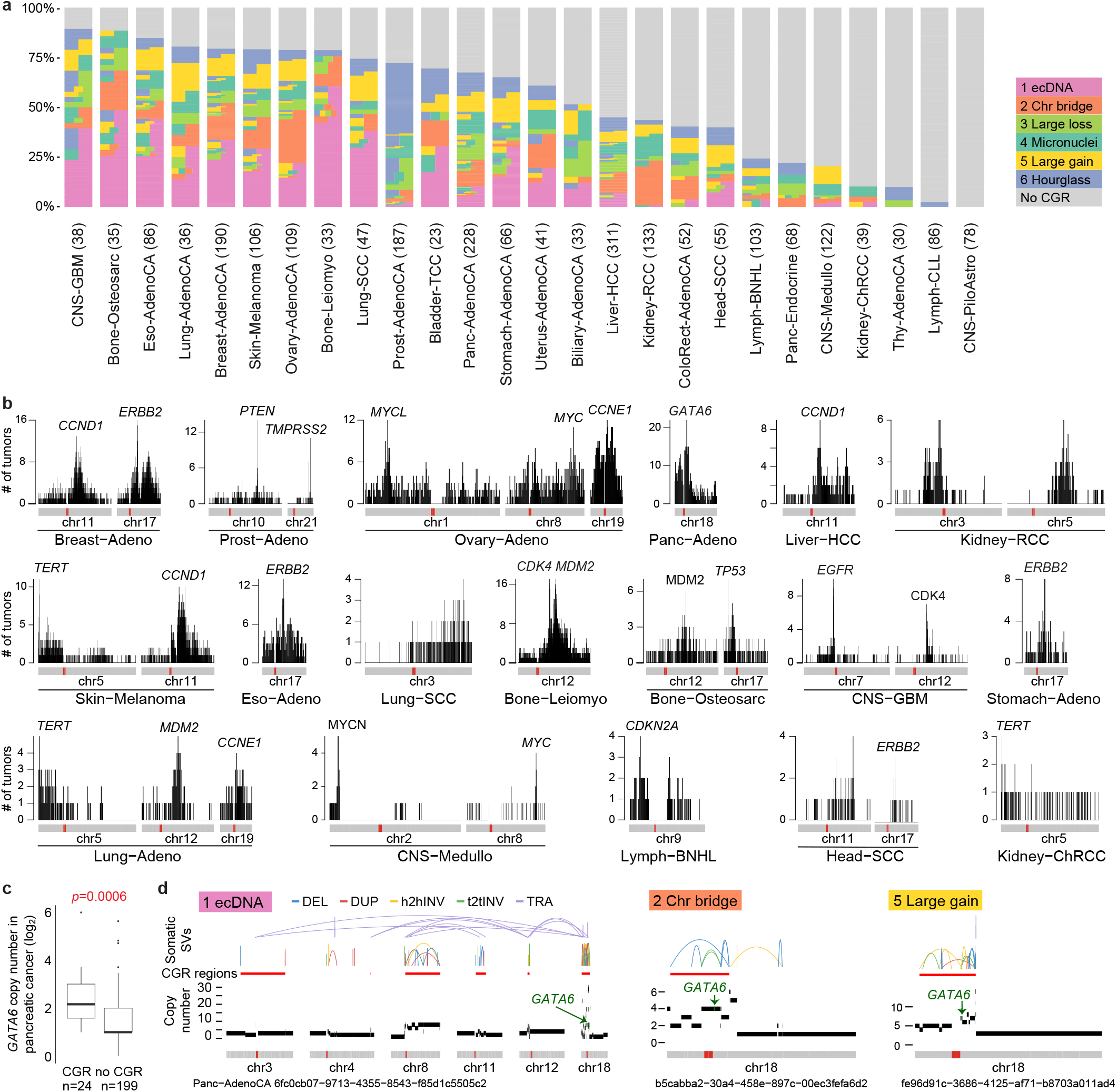
Distribution of CGRs. **a**, Percentages of CGRs in tumor types with at least 20 tumors. Tumors with more than one CGR signatures are painted by more than one colors horizontally. **b**, CGR breakpoint hotspots and cancer-driving genes. Distributions of CGR frequencies on chromosomes are shown for 18 tumor types. Each vertical line represents the number of tumors with CGR breakpoints in a 100 kb window. **c**, *GATA6* frequently amplified by CGRs in pancreatic cancers. *P* value is calculated by Wilcoxon rank sum test. **d**, Examples of *GATA6* amplified by CGRs of different signatures in pancreatic cancers. Locations of *GATA6* in three CGR regions are shown by green arrows.

CGRs also have uneven distribution across the genome (Supplementary Figure S4b). It was reported that regions with frequent CGRs often carry major cancer-driving genes, such as *ERBB2* in breast cancer^32^, *EGFR* in glioblastoma^3^ and *TERT* in chromophobe renal cell carcinoma^33^. In fact, we were able to find cancer-driving genes within 3Mb of the majority of the CGR hotspots including *CCND1*, *ERBB2*, *PTEN*, *TMPRSS2*, *MYCL*, *MYC*, *CCNE1*, *GATA6*, *TERT*, *CDK4*, *MDM2*, *TP53*, *EGFR*, *MYCN* and *CNKN2A* (Figure 2b). *GATA6* is known to be the most frequently amplified gene in pancreatic cancer^34^. We found that most of the amplifications (10.8%) are due to CGRs (Figure 2c) of different signatures (Figure 2d). Prostate cancers also have two hotspots on chromosome 21 corresponding to *TMPRSS2*-*ERG* fusions (Supplementary Figure S5) even though these fusions are usually caused by simple deletions^35^. Given the abundance of CGRs in cancers and the frequent involvement of major cancer-driving genes, we conclude that CGRs are major players of tumorigenesis.

### Genetic and clinical associations of CGRs

To better understand the mechanisms of CGR formation, we sought to identify genetic alterations associated with CGR signatures. It has been shown that *TP53* mutations are associated with chromothripsis in tumors^8,36–38^. In cell line models, *TP53* had to be inactivated so that the cells can tolerate chromothripsis without undergoing apoptosis^18–20^. After testing somatic mutations in all protein-coding genes that are mutated in at least 10 tumors in PCAWG for each CGR signature (except Signature 6), we observed Signatures 1, 2, 4 and 5 are significantly associated with *TP53* mutations (Figure 3a) with FDRs of 2.5e-10, 1.9e-3, 2.0e-4 and 4.54e-10 respectively. Interestingly, Signatures 3 and 6 (in an extended prostate cancer cohort) are significantly associated (FDRs 1.4e-2 and 5.9e-2 respectively) with mutations in Speckle Type BTB/POZ Protein (*SPOP*) (Figure 3a), a subunit of an E3 ubiquitin ligase complex involved in protein ubiquitination and degradation. We will study Signature 6 in a greater detail in a later section.

**Figure 3.**
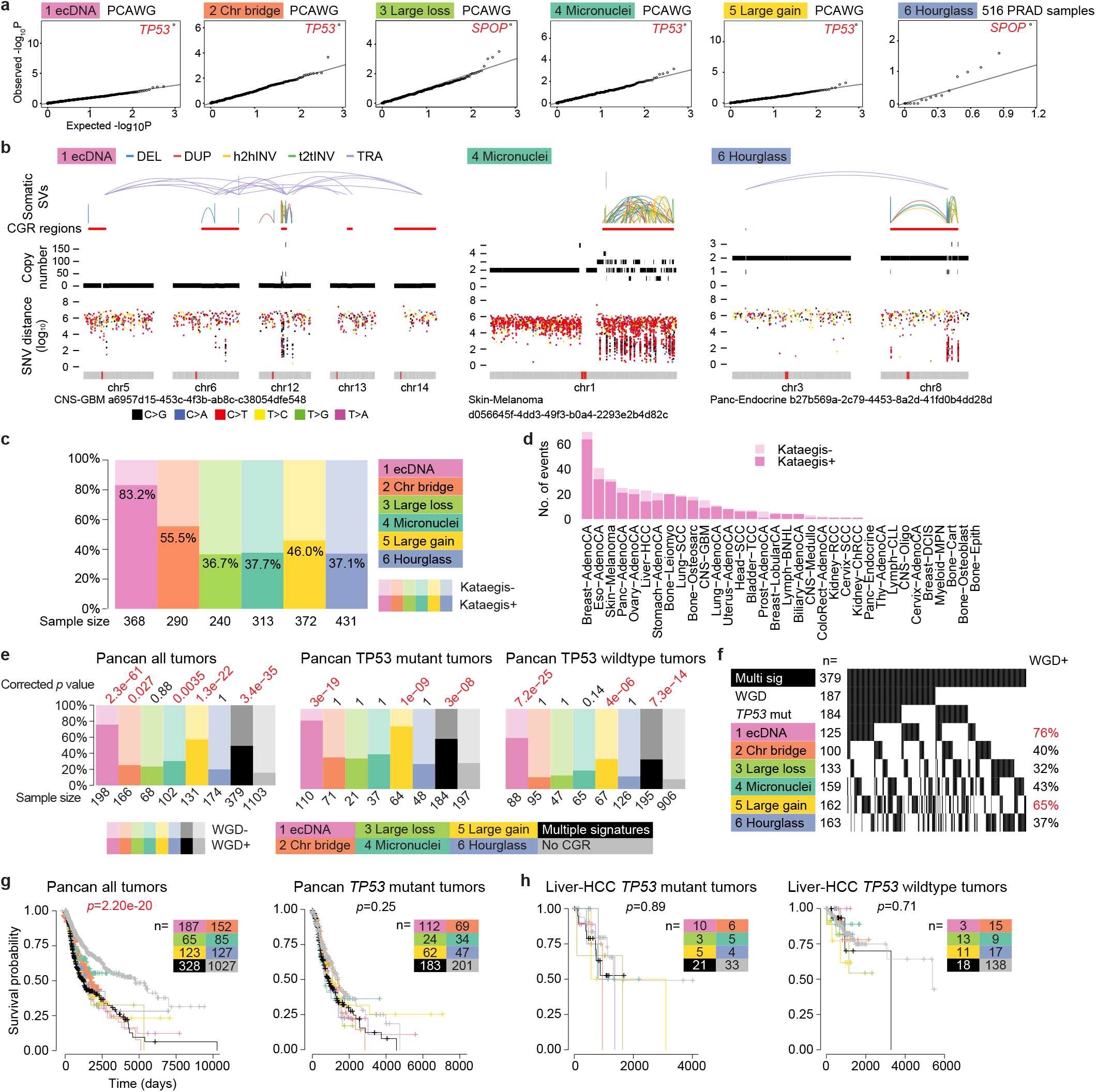
Genetic and clinical associations of CGRs. **a**, Q-Q plots of statistical associations between mutations in protein-coding genes and CGR signatures. Each dot represents a gene. Observed and expected *p* values are calculated by Fisher’s exact tests and permutation tests. **b**, Examples of kataegis co-occurring with CGRs. The top and middle panels are SV and copy number profiles of CGRs. The bottom panels are inter-mutational distances. Low distances indicate clustered SNVs. **c**, Percentages of CGRs with co-occurring kataegis. **d**, Frequencies of Signature 1 CGRs with and without kataegis in different tumor types. **e**, Associations of CGR signatures with WGD. *P* values are calculated by Fisher’s exact tests with Bonferroni correction. **f**, WGD frequencies in tumors with multiple CGR signatures. **g**, Associations between patient survival and CGR signatures with all tumor types combined. **h**, Associations between patient survival and CGR signatures in liver cancer. Log-rank test is used to calculate *p* values in **g** and **h**.

Kataegis, clustered somatic SNVs, is known to be associated with chromothripsis^4,39^. We found that kataegis events co-occur with CGRs (Figure 3b) in 1,004 out of 2,014 cases (50%). In particular, 83% of Signature 1 CGRs are accompanied by kataegis in contrast to around 40% in other signatures (Figure 3c). Such enrichment is present in almost all tumor types (Figure 3d). Interestingly, in melanoma, kataegis co-occurs with the vast majority of CGRs in all six signatures (Figure 3d, Supplementary Figure S6).

Aneuploidy is known to promote genome instability and chromothripsis^8,40–42^. We then asked if any CGR signatures are associated with whole genome duplication (WGD). When all tumors are considered, most CGR signatures are associated with WGD (Figure 3e). However, when controlled for *TP53* mutation status, only tumors with Signatures 1 and 5 as well as tumors with more than one CGR signatures are significantly associated with WGD (Figure 3e). Among tumors with multiple CGR signatures, the ones harboring Signature 1 or 5 CGRs are more likely to harbor WGD (Figure 3f). We further investigated tumor-type-specific effects. Although sample sizes are limited, Signatures 1 and 5 remain significantly associated with WGD in several tumor types such as ovarian, pancreatic, stomach cancers and melanoma (Supplementary Figure S7). In summary, two CGR signatures, Signatures 1 and 5, are associated with WGD, while other signatures are not.

Chromothripsis has been linked to poor patient survival^8,26,43^. When all patients in PCAWG cohort are tested together, CGRs are associated with poorer survival (Figure 3g left panel). However, it is possible that the poor outcome in these patients is because of impaired cell cycle checkpoints in the tumors (e.g. caused by *TP53* mutations). When controlled for *TP53* mutation status, patients with or without CGRs in their tumors have comparable survival (Figure 3g right panel). Similarly, CGR status did not predict survival when both tumor types and *TP53* mutation status are controlled for (Figure 3h and Supplementary Figure S8). Note that, our results do not conflict with previous reports showing that ecDNA is associated with poor patient outcome^25^, because ecDNA can arise from both simple genomic rearrangements and CGRs.

### CGR breakpoint biases

The uneven distribution of somatic rearrangements across the genome may be caused by biases in their formation and the breakpoint locations may provide clues about their forming mechanisms. We studied the genomic properties of CGR breakpoints by comparing observed breakpoint locations against randomly shuffled locations similar to a previous study of simple SVs^17^. All CGRs are enriched in high GC content, high gene density and early-replicated regions (Figure 4a) similar to most simple SVs^17^ suggesting that CGR may be more likely to form in open chromatin regions. The CGR breakpoints being closer to repetitive elements than expected (Figure 4a) indicates that repetitive elements may play a role in DNA fragmentation and/or ligation during CGR formation. Interestingly, CGRs of Signatures 1 and 2 tend to occur far away from telomeres while CGRs of Signatures 3, 4, 5 and 6 preferentially occur near telomeres (Figure 4a). It is possible that acentric DNA fragments resulting from breaks near the telomeres may be more likely to produce micronuclei and chromothripsis.

**Figure 4.**
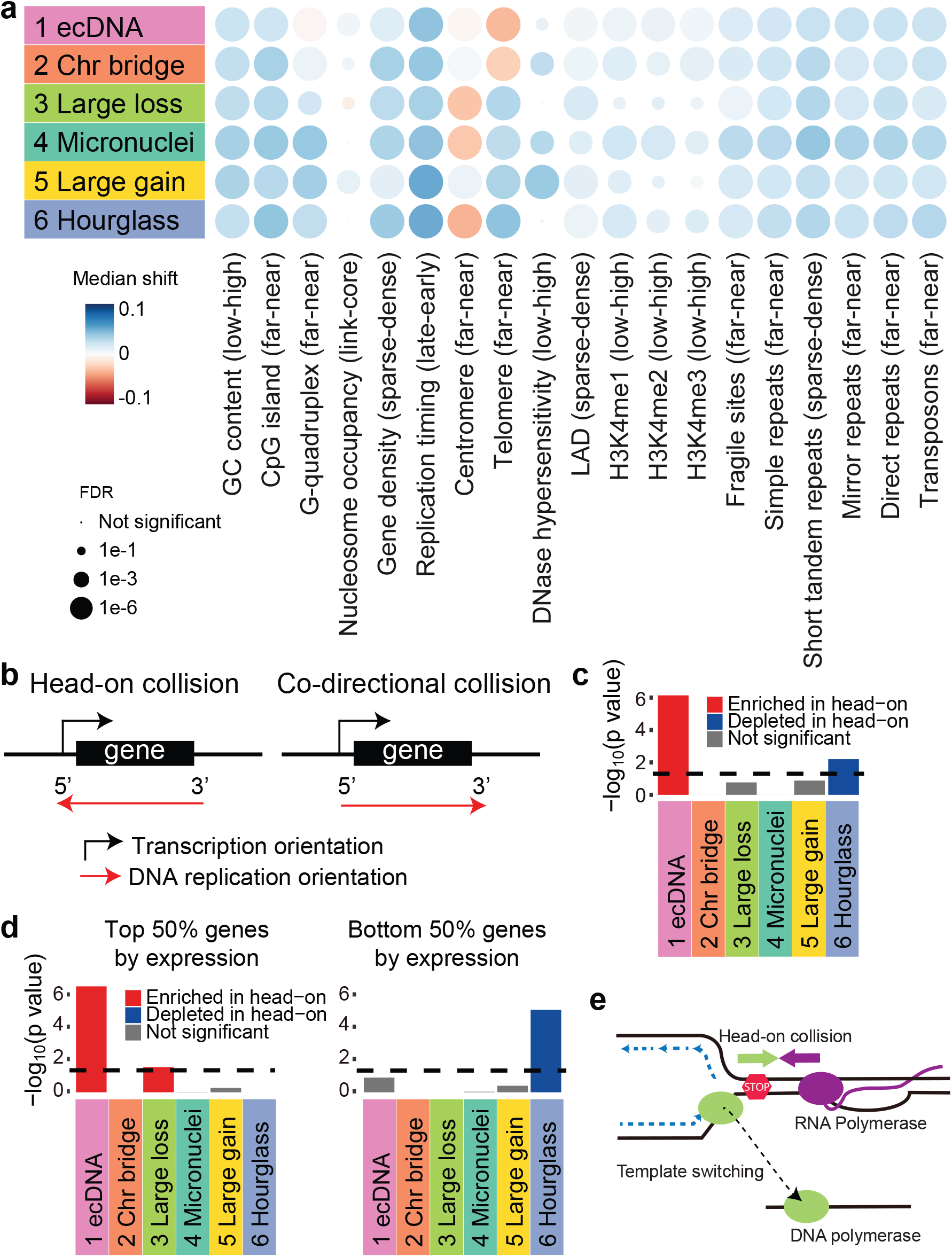
Biases of CGR breakpoints. **a**, Associations of CGR breakpoints with various genomic properties. Median shift represents the difference between the observed and random breakpoints in quantile distribution of the genomic property values. **b**, Scheme of transcription-replication collision. Head-on collision regions are defined as transcription and DNA replication orientations to be the opposite and co-directional collision regions are defined as transcription and DNA replication in the same orientation. **c**, Biases of CGR breakpoints in head-on collision regions. Bonferroni-corrected *p* values are calculated by comparing CGR breakpoints and randomly shuffled breakpoints in head-on/co-directional collision regions using Chi-square tests. **d**, Biases of CGR breakpoints in head-on collision regions controlled for gene expression level. Dashed lines in **c** and **d** denote 0.05 *p* value cutoff. **e**, Model of DNA polymerase template switching after transcription-replication collision. When DNA polymerase complex collides with RNA polymerase complex, replication fork collapses, DNA polymerase switches template and continues replication. This process may be involved in ecDNA formation.

### Role of transcription-replication collision

DNA replication stress is a major source of genome instability^44,45^. Collision between transcription and DNA replication machineries is unavoidable because both processes use the same DNA template, and can result in replication fork collapse and genome instability^46^. Some very large genes, known as common fragile sites, are hotspots for deletions due to transcription-replication collision^47^. Recently, it was reported that deletions, insertions and point mutations can frequently form when such collision is induced in a bacteria system^48^. Here, we sought to evaluate whether transcription-replication conflict contributes to the CGRs in cancer. First, we defined replication orientation based on RepliSeq data from cell line Bg02es (derived from human embryonic stem cells). The left and right replicated regions were defined as previously described^49^ (Supplementary Figure S9a) and are independent of the selection of cell types (Supplementary Figure S9b). Then, head-on and co-directional collision regions could be defined based on replication and transcription orientations (Figure 4b). We found Signature 1 breakpoints are significantly enriched in head-on collision regions (Figure 4c, Chi-square tests with Bonferroni correction) compared to randomly shuffled breakpoints. If the rearrangements are caused by transcription-replication conflict, we expect the enrichment to be dependent on gene expression. When controlled for gene expression level, we indeed found the enrichment is only significant in top 50% of the genes (highly or moderately expressed) ranked by expression level in tumors, but not in the bottom 50% (lowly expressed or not expressed) (Figure 4d). To rule out the possibility that the high gene expression being the consequence of CGRs, we performed the same test using gene expression in normal tissues and observed a similar bias (Supplementary Figure S9c) towards expressed genes. To further rule out the effect of selection, we removed breakpoints within 3 Mb of hotspots (Figure 2b) and the bias could still be observed (Supplementary Figure S9d). Previous studies based on *in vitro* experiments in cell lines reported that ecDNA can form via chromothripsis^18,23^. Our results indicated that conflicts between DNA replication and transcription may contribute to ecDNA formation in tumor tissue. When a replication fork collapses, DNA polymerase can switch to a new template and different types of genomic rearrangements can form depending on the destination of the polymerase^50,51^. Template switching upon transcription-replication collision (Figure 4e) can be a plausible mechanism to produce a circular molecule. Further studies are needed to elucidate the role of transcription-replication collision in ecDNA formation.

### Hourglass chromothripsis in prostate cancer

The Signature 6 hourglass chromothripsis is dominant in prostate cancer (Figure 2a). Chromoplexy is another form of CGR enriched in prostate cancer, lymphoid malignancies and thyroid cancer^4,11^. It is considered to be the result of ligation of simultaneously broken DNA ends of several chromosomes^11^—a complex form of reciprocal translocations^4^. We sought to address whether hourglass chromothripsis is equivalent to chromoplexy. Using two strategies to detect chromoplexy: ChainFinder^11^ and junction patterns^17^, we found hourglass chromothripsis events have little overlap with chromoplexy (Supplementary Figure S10a). In addition, most hourglass chromothripsis cases only involve one or two chromosomes (Figure 1e) while chromoplexy usually involves multiple chromosomes^11^. Other than prostate cancer, hourglass chromothripsis is commonly seen in glioblastoma and bladder cancer (Figure 2a), while chromoplexy is enriched in thyroid cancer and lymphoid malignancies^4^. Therefore, we conclude that hourglass chromothripsis is a unique type of CGRs and distinct from chromoplexy.

To test whether hourglass chromothripsis is a one-time catastrophic event, we utilized linked-read sequencing data of 23 prostate cancers^52^. We identified 10 hourglass chromothripsis events in 15 tumors including two events in an *SPOP* mutant tumor 01115468-TA3. In this tumor, one of the hourglass chromothripsis occurred on the long arm of chromosome 8 (Figure 5a). Once the rearranged tumor chromosome was reconstructed by following individual somatic SVs (Figure 5b), all rearranged DNA fragments in Figure 5b could be phased into a single haplotype using linked-read barcodes. The same tumor harbored another more complex hourglass chromothripsis involving five chromosomes (Figure 5c). Using the same procedure, we identified seven phased blocks with more than one somatic SV. There are a total of 155 somatic SVs in these seven phased blocks. If this hourglass chromothripsis results from simple SVs accumulated over time, we expect the somatic SVs to be evenly represented in two haplotypes. However, 144 of them can be phased to one haplotype which is extremely unlikely to occur by chance (*p*=1.3e-28, binomial tests, *p* values combined with Fisher’s method). These results suggest that hourglass chromothripsis events are indeed one-time catastrophic events.

**Figure 5.**
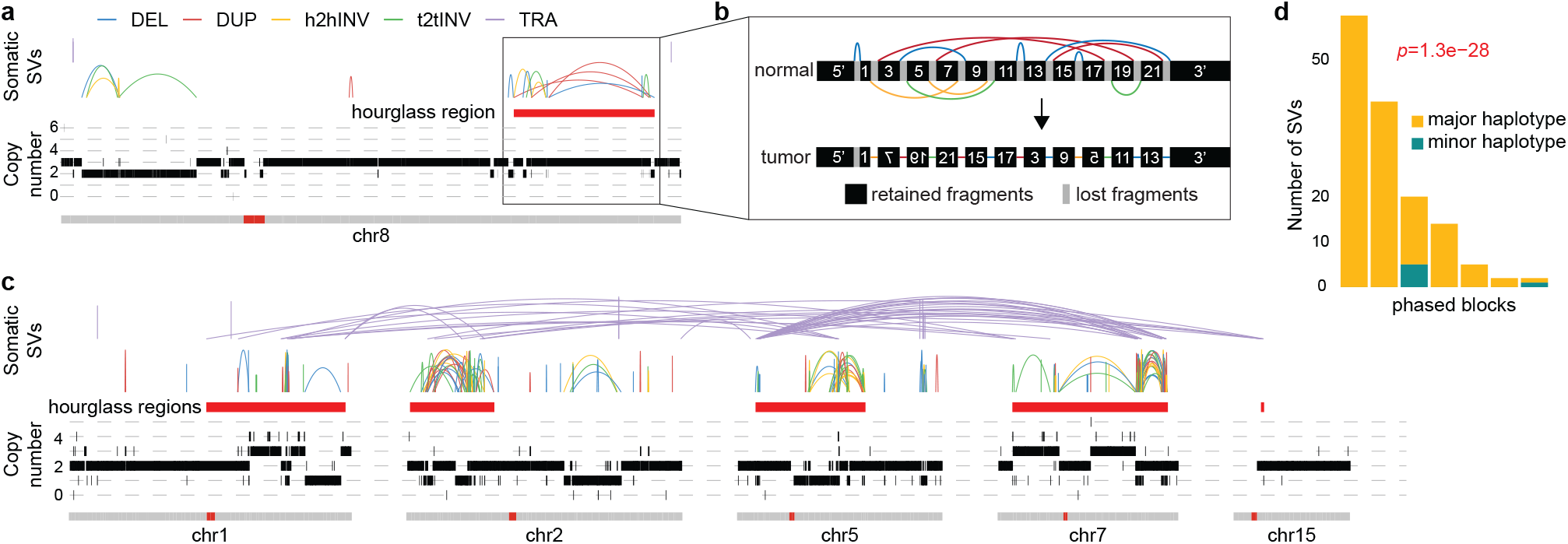
Reconstruction of hourglass chromothripsis using linked-read sequencing. **a**, An hourglass chromothripsis on chromosome 8 in a prostate cancer with *SPOP* mutation. **b**, Reconstruction of hourglass chromothripsis in **a**. Genomic segments are re-scaled for the chromothripsis region. The normal chromosome and SVs are shown on the top and the reconstructed tumor chromosome is shown at the bottom. Segments with flipped texts in tumor chromosome represent inverted DNA fragments. **c**, Another hourglass chromothripsis in the same tumor involving five chromosomes. **d**, Phasing somatic SVs in **c** using barcoded reads. Seven phased blocks with more than one somatic SVs are shown. *P* value was calculated by binomial test in each phased block and combined by Fisher’s method.

We then took advantage of additional 329 publicly available WGS prostate cancers from International Cancer Genome Consortium^53,54^ and identified another 359 CGRs (Supplementary Table S3). In the combined cohort of 516 prostate cancers, we found that mutations in *SPOP* are significantly associated with hourglass chromothripsis (*p*=3.4e-3, Fisher’s exact test, Figure 6a). *SPOP* is known to be recurrently mutated in prostate cancer and the mutations are mutually exclusive with ETS fusions (*TMPRSS2*-*ERG*, -*ETV1*, -*ETV4* and -*ETV5*)^55^. All mutations are missense mutations in the meprin and TRAF homology (MATH) domain (Supplementary Figure S10b) and potentially disrupt *SPOP*’s target binding^56^. Mutant *SPOP* is associated with micronuclei formation^57^ which suggests *SPOP* is likely to be directly involved in hourglass chromothripsis formation. Signatures 3 and 4 are also associated with *SPOP* mutations (*p*=1.4e-10 and 1.5e-3, respectively, Fisher’s exact test, Figure 6a). In addition, *SPOP* mutations are associated with simple genomic rearrangements as well (Supplementary Figure S10c). Based on these results, we conclude that SPOP is likely to be a gatekeeper of genome stability similar to p53. Mutant SPOP may allow the cells to tolerate various types of genomic rearrangements.

**Figure 6.**
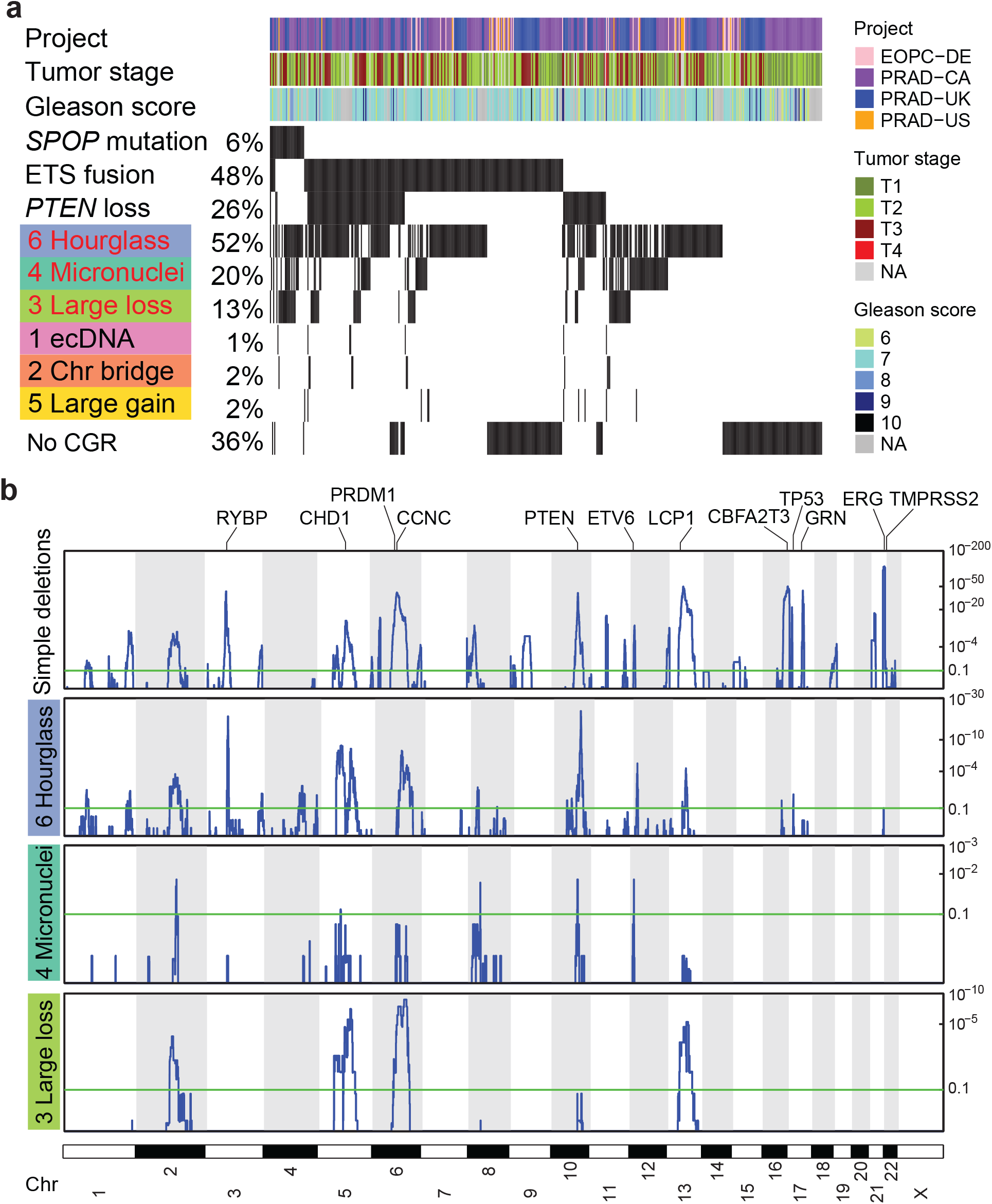
Hourglass chromothripsis in prostate cancer. **a**, CGRs and genetic alterations in a combined cohort of 516 prostate cancers. Red texts depict CGR signatures significantly associated with *SPOP* mutations. **b**, Recurrently deleted regions in four types of copy-loss-associated rearrangements in prostate cancer.

Next, we sought to investigate the functional consequences of hourglass chromothripsis in prostate cancer. We identified recurrently deleted regions resulting from Signatures 3, 4 and 6 as well as simple deletions using GISTIC^58^. We found that CGRs, especially hourglass chromothripsis events, mostly delete the same regions as those regions deleted by simple deletions leading to loss of cancer driver genes such as *PTEN* (Figure 6b). This suggests that CGRs are under positive selection similar to recurrent simple deletions. A few GISTIC peaks of simple deletions are not found in CGRs including the most frequent one in chromosome 21q22 (Figure 6b) causing *TMPRSS2*-*ERG* fusions^35^. *TMPRSS2*-*ERG* fusions should be less likely to result from CGRs because both *TMPRSS2* and *ERG* reside on chromosome 21 and are 3 Mb away from each other. The easiest way to form a fusion gene is to connect the two genes by a simple deletion. The chance of forming fusion gene through CGRs is expected to be much lower since DNA fragments in CGRs are randomly ligated. Nonetheless, we still observed 11 *TMPRSS2*-*ERG* fusions resulting from hourglass chromothripsis events in 187 prostate cancers in the PCAWG cohort (Figure 2b and Supplementary Figure S5), which further suggested that hourglass chromothripsis is a major player of tumorigenesis in prostate cancer and the impact of CGRs is similar to that of simple SVs.

## Discussion

In this study, we used an unbiased approach to deconvolute CGR signatures based on the event topology. Our strategy does not rely on any prior knowledge of CGR mechanisms and the benchmarking results demonstrate high concordance with experimentally induced CGRs (Figure 1d). Our approach has several limitations. First, the CGR signatures were decomposed based on five genomic features which might not adequately reflect all the variations in topology of different molecular mechanisms. It is possible that there are other informative features we did not model. In addition, most CGR signatures have one dominant feature (Figure 1b). So, some CGR events might be misclassified. For example, Signature 2 has the highest score in telomere loss (Figure 1b) which is consistent with chromatin bridge mechanism^20^. If the breakpoint junction is too close to the telomere, the event would have low telomere-loss score and be misclassified as a different signature. Indeed, a small fraction of CGR events are misclassified in benchmarking cases (Figure 1d). Nevertheless, we were able to correctly classify most of the CGR events from several independent experimental studies (Figure 1d) suggesting the good overall performance of Starfish. Another limitation of our study is sample availability. Although PCAWG is the largest pan-cancer WGS dataset to date, most tumor types have less than 100 samples (Figure 2a). Most subtypes are not adequately represented. For example, there are over 100 subtypes of soft-tissue sarcomas^59^, whereas only a handful of them were profiled in PCAWG. Furthermore, the majority of the patients are Caucasians. Tumor-type specific, subtype specific and ethnic-group specific CGRs may be missed in the current study.

Although we found that most CGR hotspots are associated with cancer-driving genes (Figure 2b), there are exceptions. In clear cell renal cell carcinoma, chromosomes 3 and 5 are known to harbor very frequent chromothripsis events which lead to 3p loss and 5q gain^31^. No driver gene is found near the hotspots on either chromosome. This might be because these tumors are driven by losses and gains of multiple genes^60,61^ on 3p and 5q including *VHL*, *PBRM1*, *SETD2*, *BAP1* and *SQSTM1*.

In a recent study^23^, when certain cell lines were exposed to drugs, various types of genomic alterations could be observed such as arm-level copy gains, simple rearrangements, BFB cycles, chromothripsis and ecDNA with various configurations. The experimental results demonstrated the exceeding complexity of genomic alterations. What we observe here in human tumors is highly consistent with the experimental observation that one gene could be amplified through several distinct mechanisms (Figure 2d). In our previous study, we have reported that *EGFR* in glioblastoma could be amplified as a single fragment in DM or co-amplified with many other fragments^3^. In many Signature 1 CGRs, some fragments in the CGR regions (e.g., chromosomes 1, 12 and X in Figure 1c ecDNA of leiomyosarcoma) are amplified while others are not amplified or amplified at a low level (e.g., chromosomes 6 and 7 in Figure 1c ecDNA of leiomyosarcoma). Our results also revealed unique features of ecDNA—their breakpoints are enriched in transcription and DNA replication head-on collision regions (Figure 4c and 4d). This suggests that although ecDNA can form through micronuclei-induced and chromatin-bridge-induced chromothripsis events *in vitro*^18,23^, transcription activity and replication stress may play important roles in ecDNA formation during tumorigenesis.

It is possible that hourglass chromothripsis is a special case of micronuclei-induced chromothripsis with less amount of DNA loss because both display small number of copy states and oscillating copy numbers. If both events are the results of random ligation of shattered DNA fragments, individual fragments being retained or lost should follow a Bernoulli distribution. In hourglass chromothripsis events, large fragments are retained while small ones are lost. If they have the same forming mechanism as micronuclei-induced chromothripsis, hourglass chromothripsis should be rare. However, hourglass chromothripsis is more common than micronuclei-induced chromothripsis (Figure 1b) which suggests they are unlikely to form via random chromosomal shattering and ligation. In addition, other evidences also suggest that these two types of chromothripsis events are distinct. The most dominant feature of hourglass chromothripsis is high breakpoint dispersion score while micronuclei-induced chromothripsis has the lowest score (Figure 1b). Hourglass chromothripsis is abundant in prostate cancer while micronuclei-induced chromothripsis is not enriched in any tumor types (Figure 2a). Intriguingly, the breakpoints of hourglass chromothripsis, but not micronuclei-induced chromothripsis, are depleted in transcription-replication head-on collision regions (Figure 4c) which is the opposite of ecDNA. Such depletion depends on gene expression that the depletion is only observed in bottom 50% of the genes ranked by expression level but not in the top 50%, which is again the opposite from ecDNA (Figure 4d). These results indicate that the formation of hourglass chromothripsis might be associated with transcription-coupled repair. These notable differences between hourglass and micronuclei-induced chromothripsis suggest that hourglass chromothripsis has a distinct mechanism of formation. Further studies using experimental approaches will be necessary to reveal the detailed molecular mechanisms.

## Methods

See Online Methods section.

## Acknowledgement

This work was supported by NCI grant K22CA193848 (L.Y.), University of Chicago and UChicago Comprehensive Cancer Center. We thank the Center for Research Informatics at the University of Chicago for providing the computing infrastructure and Matthew Stephens, Xin He and Michelle Le Beau for helpful suggestions.

