## Supplementary Methods for "Mutational signatures of complex genomic rearrangements in human cancer"

**Samples and variant calling**

PCAWG WGS data for 2,428 tumor and matched-normal pairs across 37 cancer types^1^ were used for the majority of this study unless otherwise noted. All variants (SNVs, CNVs, SVs, tumor purity, ploidy and WGD) were called by the PCAWG consortium using multiple algorithms^2^. Kataegis was detected by SeqKat^3^. Another set of 329 WGS prostate cancers from ICGC were used to study CGRs in prostate cancer. Somatic variants were downloaded from ICGC data portal (<https://dcc.icgc.org/>). All genomic coordinates were based on the hg19 reference genome.

**Identification of CGRs**

Modified ShatterSeek^1^ was used to identify CGR “seed” regions based on interleaved SVs, goodness-of-fit, fragment joins test, chromosomal enrichment test, and exponential distribution of breakpoints test. Oscillating-copy-state criteria was removed from the original ShatterSeek package. In each sample, linked regions were identified if they were connected by at least two translocations within 10 kb of any seed regions. The search was performed iteratively until no new linked regions could be found. Finally, a CGR event was defined as all connected seed and linked regions combined (Supplementary Figure S11). The somatic SVs of CGR were defined as all inter-chromosomal translocations and interleaved intra-chromosomal SVs in the CGR regions. The remaining SVs were defined as simple SVs.

**CGR signature deconvolution**

Twelve features were initially selected to comprehensively describe CGR topology of 2,014 events including breakpoint dispersion score, copy loss percentage, copy loss density (number of copy loss fragments per 10Mb, log scale), copy gain percentage, copy gain density (number of copy gain fragments per 10Mb, log scale), number of copy states (log scale), median copy number change (log scale), maximum copy number (log scale), highest telomere loss percentage, ratio of telomere loss and CGR loss sizes (log scale), median breakpoint microhomology (log scale), and median breakpoint insertion size (log scale). In Figure 1a, the size of CGR region [s] equals to $\sum a:g$. Breakpoint dispersion score was defined as mean absolute deviation of [a:g] which measured the randomness of breakpoint distribution of a CGR. Copy loss and copy gain percentages were calculated by $(\sum b,f)$/[s] and ($\sum d,g)$/[s]. Copy loss and copy gain densities were length[b,f]/([s]/10Mb) and length[d,g]/([s]/10Mb). Telomere loss percentage was t/L. To reduce collinearity among these features, the features with high correlation (abs(correlation coefficient)>0.5) were removed, including copy loss density, copy gain density, number of copy states, median copy number change, and ratio of telomere loss and CGR loss sizes. Features with small variance were further removed including median breakpoint microhomology and median breakpoint insertion size. After feature selection process, five features were kept for downstream analysis: breakpoint dispersion score, copy loss percentage, copy gain percentage, maximum copy number and highest telomere loss percentage. Then, a 2,014 x 5 matrix was constructed for all CGRs. All values were transformed into z scores. Four clustering methods (k-means based on Euclidean distance, partitioning around medoids [PAM] based on Euclidean distance, hierarchical clustering based on Pearson distance, and hierarchical clustering based on Euclidean distance) were used to perform unsupervised clustering. Optimal cluster number K and the best clustering method were determined based on four scores: Silhouette score, C-index, Calinski-Harabasz score, and Dunn-index. The K maximizing the ratio of intra-cluster similarity/inter-cluster similarity was selected. The final clustering was performed by R package “Consensuscluster Plus” using PAM with 0.9 item and 5000 iterations.

**Building neural network classifier**

The feature matrix and CGR signature labels from Figure 1b were used to construct a signature classifier. The R package “neuralnet” was used to train a one layer neural-network classifier with 5-fold cross validation using 70% of CGR events as the training data and 30% of them as the validation data for each iteration. One to sixteen neurons were used in each iteration with 10e6 steps to converge at the error of 0.01. The model with 8 neurons was selected.

**Benchmarking using experimentally induced chromothripsis**

A total of 160 WGS samples from three published studies^4–6^ were used to benchmark our CGR signatures. Somatic SVs and CNVs were obtained from the corresponding publications. The CGRs were detected as described above. The feature matrices from individual studies were combined with PCAWG feature matrix and normalized. The CGR signatures were predicted using the neural network classifier.

**Associations between somatic SNVs and CGR signatures**

Protein-coding gene mutation status and the presence/absence of CGR signatures were used to construct two by two contingency tables across all PCAWG samples. Only genes mutated in more than 10 tumors were considered. Fisher’s exact test was used to compute *p* values and false discovery rate was computed by Benjamini- Hochberg procedure for each CGR signature. Significant genes were selected by FDR<0.1. Mutation status was permuted using R package “QQperm” with 1000 iterations to calculate expected *p* values and generate Q-Q plots.

**Survival analysis**

Survival analysis of samples with different CGR signatures was performed with Kaplan-Meier method by using the R package “survival”. *TP53* mutation status and tumor type were further controlled for stratified survival analysis. *P* values were calculated by log-rank tests.

**CGR signatures in prostate cancers**

The CGR signatures were predicted in 329 ICGC prostate cancers^7,8^ and 23 prostate cancers sequenced by linked reads^9^ as described in the benchmarking section using neural network classifier.

**CGR breakpoint enrichment analysis**

Two random SVs were generated for each CGR SV by fixing the SV size and orientation and then placing them randomly to unique mappable regions of the same chromosome. CpG island annotation was downloaded from UCSC Genome Table Browser (genome.ucsc.edu/cgi-bin/hgTables). G-quadruplex (G4) clusters and fragile site positions were obtained from previous studies^10,11^. Distances to CpG islands and G4 structures were log10 transformed. Nucleosome occupancy (mean values at 5 base-pair resolution) of K562 cell line, replication timing of NHEK (derived from normal skin) cell line, DNase hypersensitivity (average imputed negative log p-value in 1 kb window) of GM12878 cell line, histone modifications (average signal values in 1 kb window) of Gm12878 cell line, and repeat sequence annotations were downloaded from UCSC Genome Table Browser. Gene density was calculated as the proportion of bases in protein-coding genes in 1 Mb window based on GENCODE v19. Lamina associated domain (LAD) (proportion of bases in 1 Mb window) in the Tig3 cell line of normal human embryonic lung fibroblasts were obtained from a previous study^12^. The density of short tandem repeats was calculated as the proportion of bases belonging to short tandem repeats in a 3 kb window. For each genomic property, a quantile distribution for the genomic property values at the observed breakpoints and randomly shuffled breakpoints were generated and the median shift was calculated as “the median quantile of observed breakpoints” minus “the median quantile of shuffled breakpoints”. A Kolmogorov-Smirnov test was conducted comparing normalized genomic property values from observed and random breakpoints, and a Benjamini-Yekutieli correction for false discovery rate was performed. All statistical analyses were performed in R 3.6.0.

**Transcription-replication collision**

Replication timing profiles for the Bg02es cell line as well as Bj, GM12878, HelaS3 and K562 were obtained from ENCODE (<https://www.encodeproject.org/>). Left- and right-replicated regions were defined as regions where the changes of replication timing were more than 0.03 per kb (Supplementary Figure 9a). In Bg02es, a total of 1.66 Gb sequences from the reference genome had defined replication orientation. The replication orientations defined from Bg02es were used for statistical tests. Transcription orientations were derived using gene annotation of GENCODE v19. Only breakpoints in protein-coding genes were considered. Breakpoints in overlapping genes with different orientations were discarded. Gene expression levels (upper-quartile-normalized fragments per kb per million mapped reads [FPKM-UQ]) of tumor tissues were available from PCAWG^13^ and gene expression levels of normal tissues (only a subset of tumor types have matched-normal tissues) from TCGA samples were downloaded from Genomic Data Commons (<https://portal.gdc.cancer.gov/>). Median expression levels of protein-coding genes within each tumor type were used. The numbers of SV breakpoints observed in tumors and the numbers of randomly generated SV breakpoints were used to test breakpoint enrichment in head-on collision regions using Chi-square test with Bonferroni correction.

**Reconstruction of hourglass chromothripsis using linked reads**

A total of 23 prostate cancer tumor/normal pairs sequenced by 10x linked reads were obtained from a recent study^9^. Somatic SNVs were called by the previous study^9^. Somatic SVs were called by Long Ranger (<https://github.com/10XGenomics/longranger>) and SvABA^14^. Copy number segments in tumors were identified by BIC-Seq^15^, and integer copy number was calculated by Sequenza^16^. Somatic CNV and SV calls were inspected by IGV and Loupe. Phased blocks were inferred in the tumor samples by Long Ranger based on heterozygous SNVs (both germline and somatic) and barcoded reads. The barcoded reads with at least three heterozygous SNVs were retrieved to assign unique haplotypes by gemtools^17^. Somatic SVs in the tumor samples were phased using barcodes and shared SNVs. Phased blocks were further merged if they shared at least ten barcoded read pairs or were connected by at least two somatic SVs. The haplotype contained more SVs was assigned as the major haplotype and the other one was assigned as the minor haplotype. Binomial test was performed in each of the seven phased blocks with more than one somatic SVs to test the enrichment of SVs in the major haplotype, and the *p* values were combined by Fisher’s method.

**Code availability**

The Starfish package is available at <https://github.com/yanglab-computationalgenomics/Starfish>.
