## Supplementary Figures for "Mutational signatures of complex genomic rearrangements in human cancer"

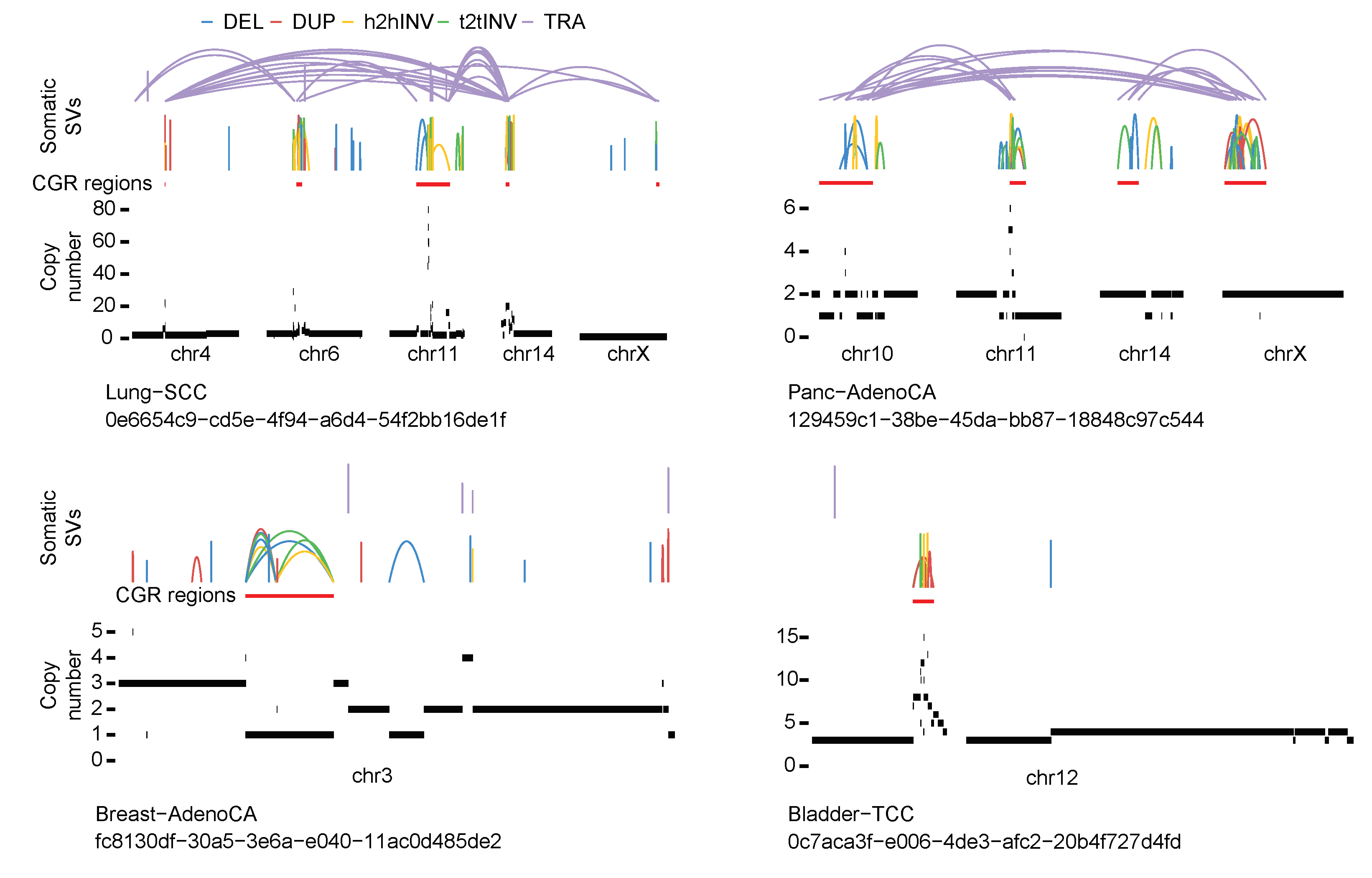

**Figure S1. Examples of CRGs not detected by the original version of ShatterSeek due to oscillating copy-number state requirement.**

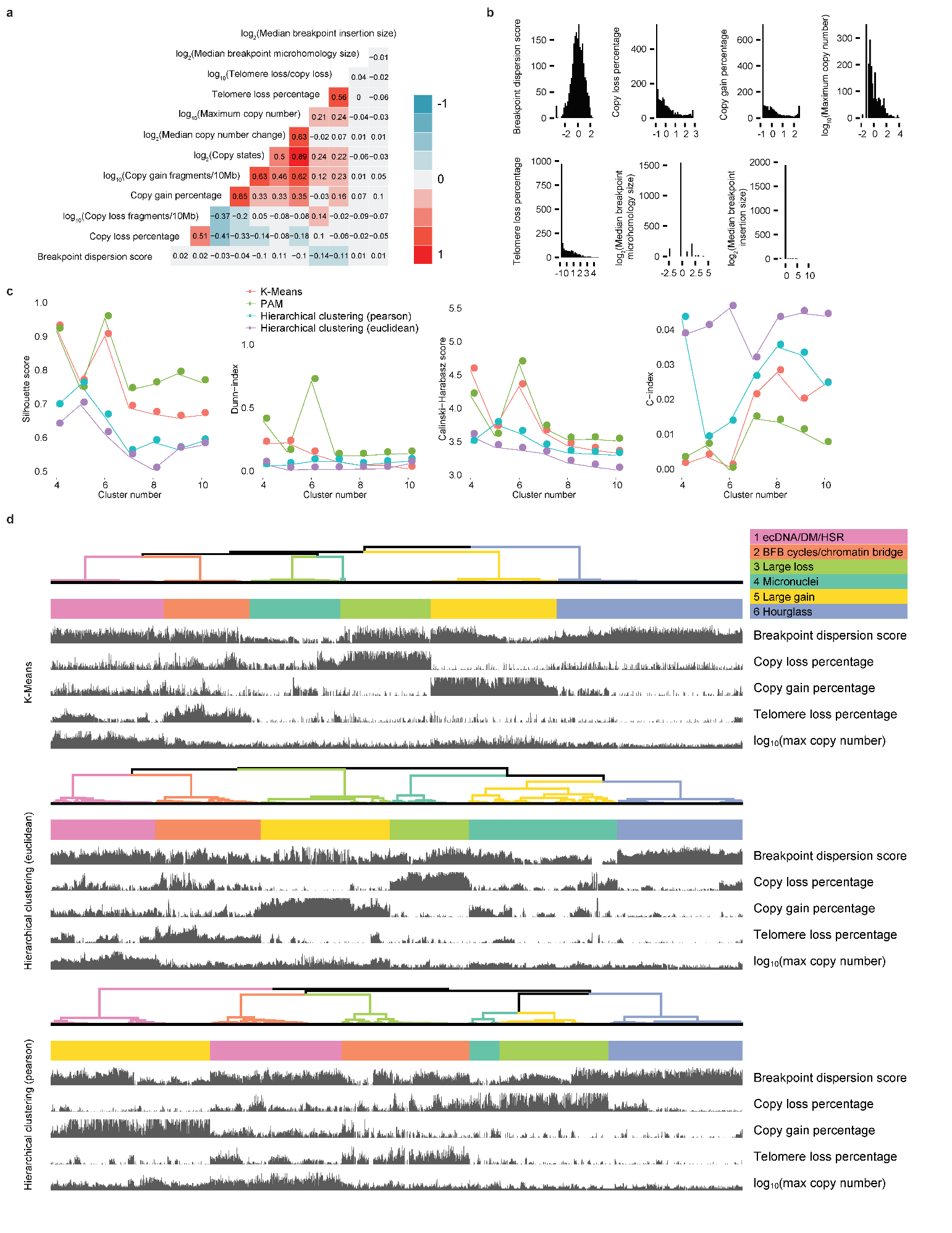

**Figure S2. Clustering of CGRs. a,** Correlation heatmap of twelve genomic features. The colors represent the correlation coefficients. **b,** Distributions of seven genomic features with low correlations. Two features (median breakpoint microhomology size and median breakpoint insertion size) have low variations and are not used in clustering. **c,** Four indexes to evaluate number of clusters and four clustering methods. **d,** Clusters produced by three other methods (K-means, Hierarchical clustering with Euclidean distance and Hierarchical clustering with Pearson distance).

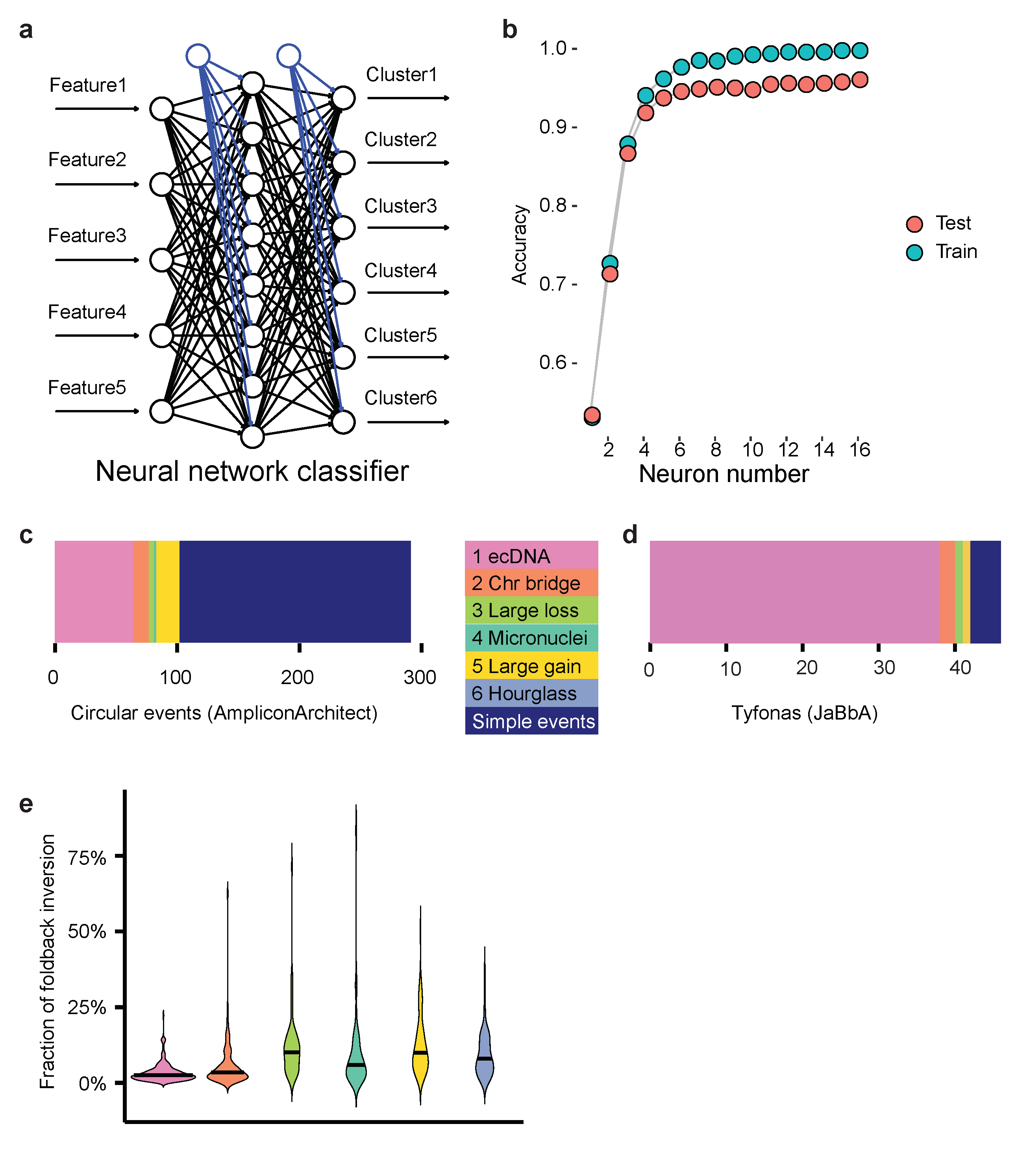

**Figure S3. CGR signature classifier and benchmarking CGR signatures. a,** Scheme of neural network classifier of CGR signatures. **b,** Testing performance of neural network classifier of CGR signatures. **c,** Comparison between circular events predicted by AmpliconArchitect and CGR signatures. **d,** Comparison between tyfonas events predicted by JaBbA and CGR signatures. **e,** Fractions of foldback inversions in six CGR signatures.

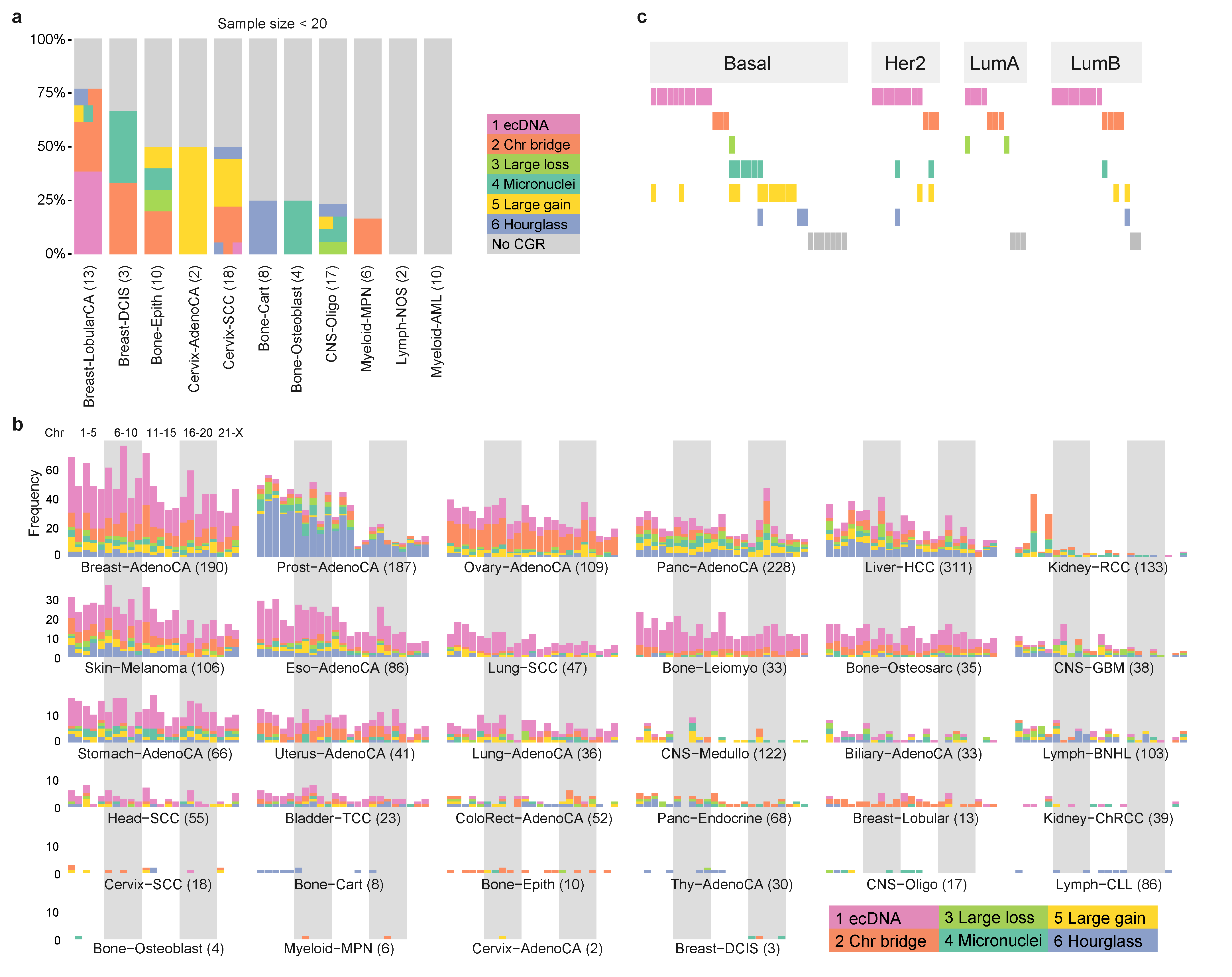

**Figure S4. Distribution of CGRs in different tumor types. a,** Percentages of CGRs in tumor types with small sample size. Tumors with more than one CGRs are painted by more than one colors **b,** Frequencies of CGRs per chromosome in different tumor types. CGRs on chromosomes 1 to X are shown with 23 bars **c,** Frequencies of CGRs in four breast cancer subtypes.

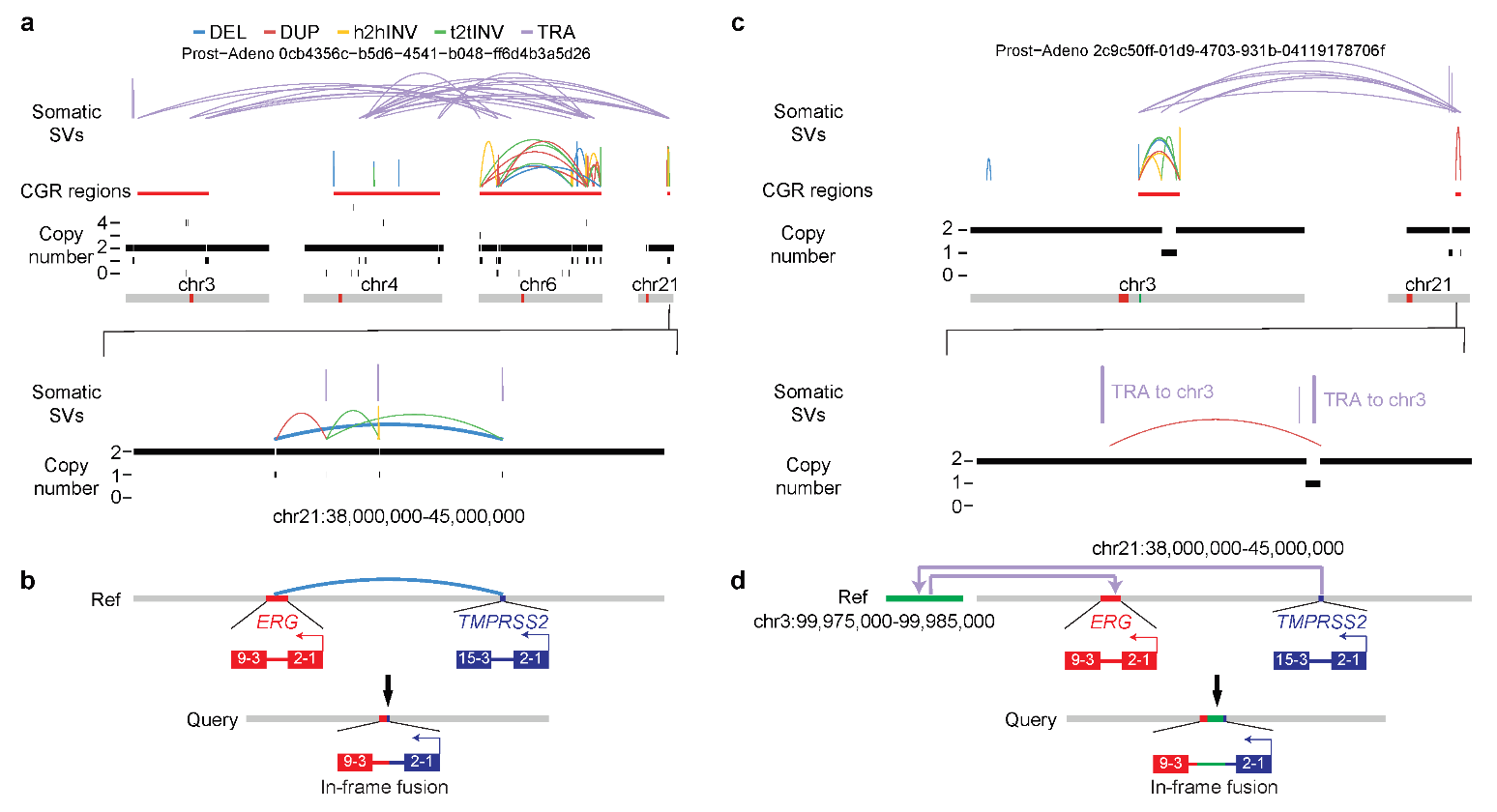

**Figure S5. *TMPRSS2*-*ERG* fusions generated by CGRs in prostate cancer. a** and **c**, SV and CNV profiles of two CGRs in prostate cancers. The lower panels show zoom-in regions of *TMPRSS2* and *ERG* on chromosome 21. Fusion producing SVs are bolded. **b** and **d**, Reconstruction of fusions generated by CGR SVs.

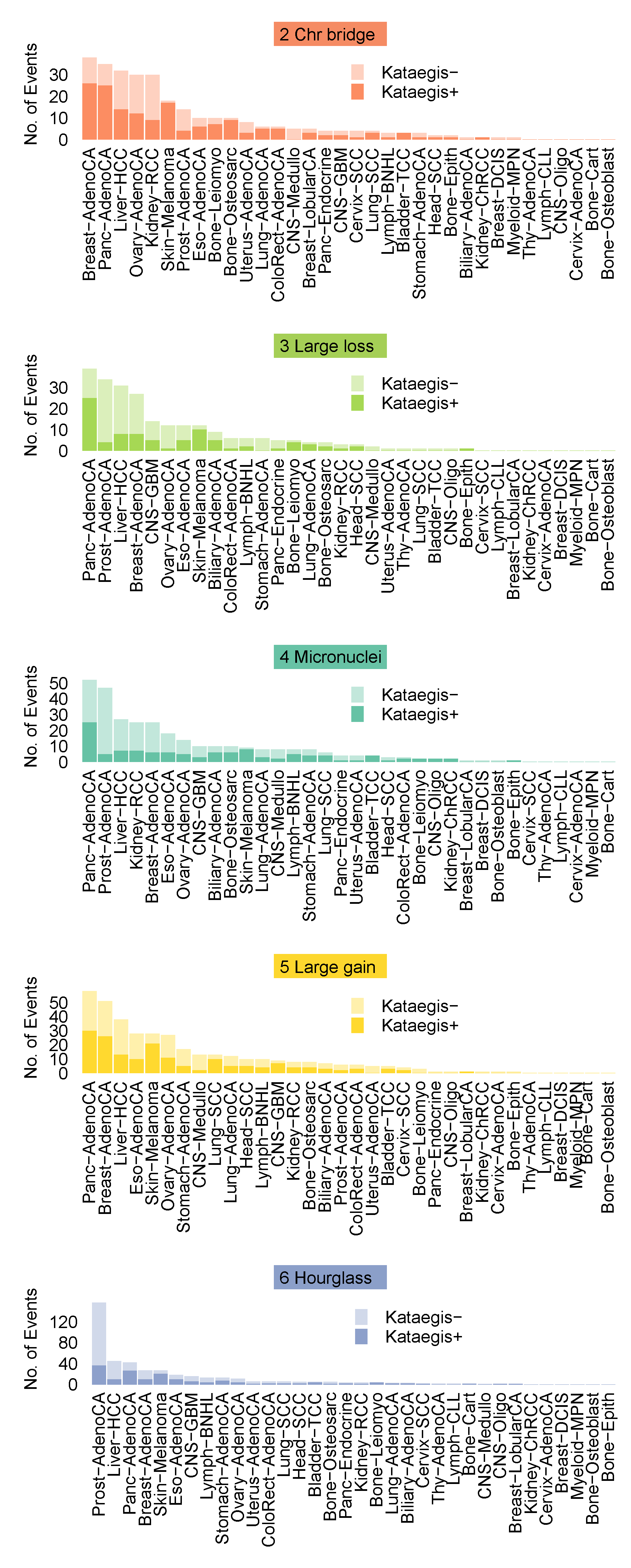

**Figure S6. The numbers of CGRs with and without co-occurring kataegis in each tumor type stratified by five CGR signatures.** Signature 1 is shown in Figure 3d.

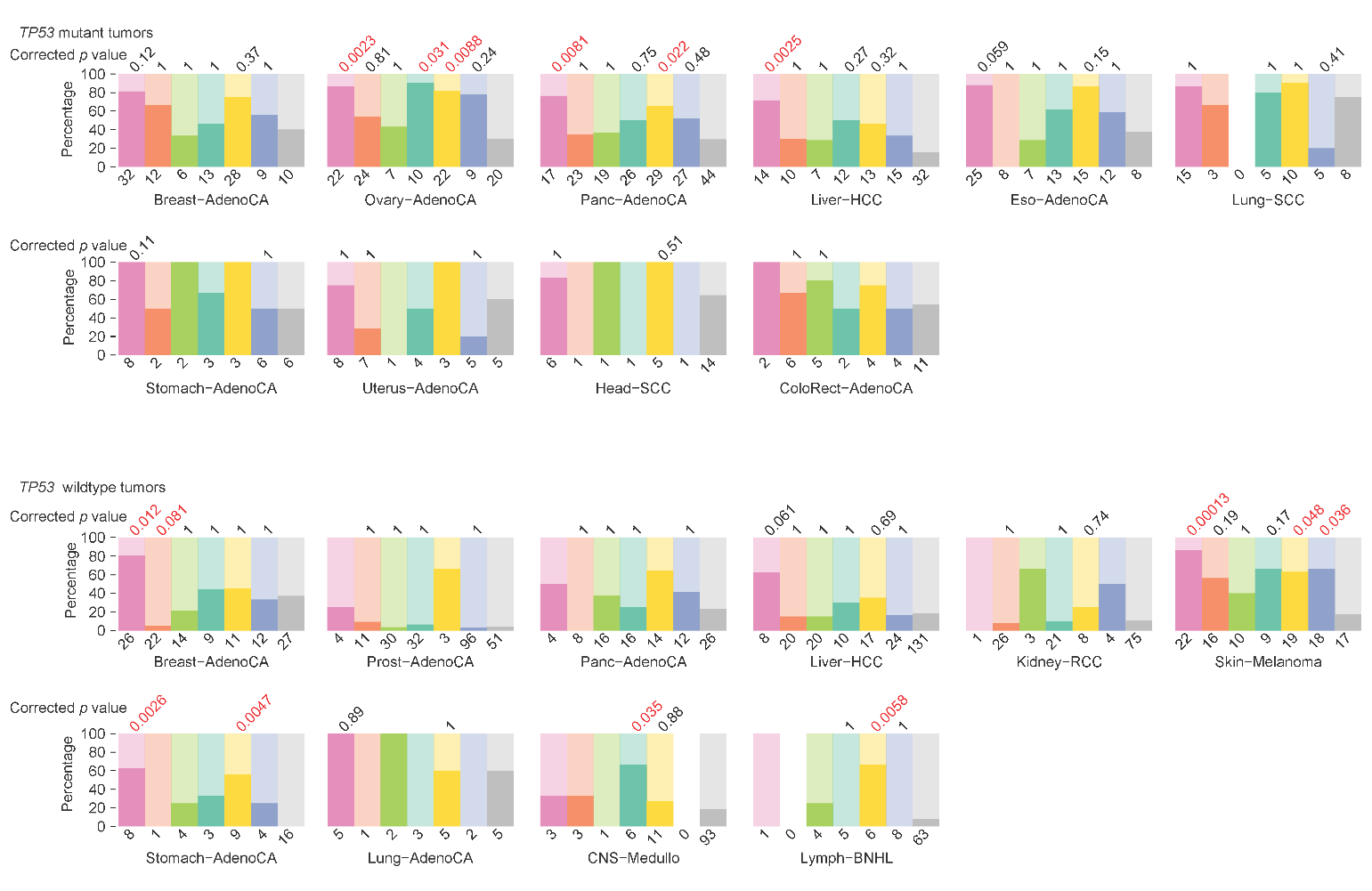

**Figure S7. Percentages of CGR-carrying tumors with and without WGD stratified by *TP53* mutation status and tumor type.** Only tumor types with at least five samples in no CGR group and at least five samples in any one of the CGR signature group are displayed.

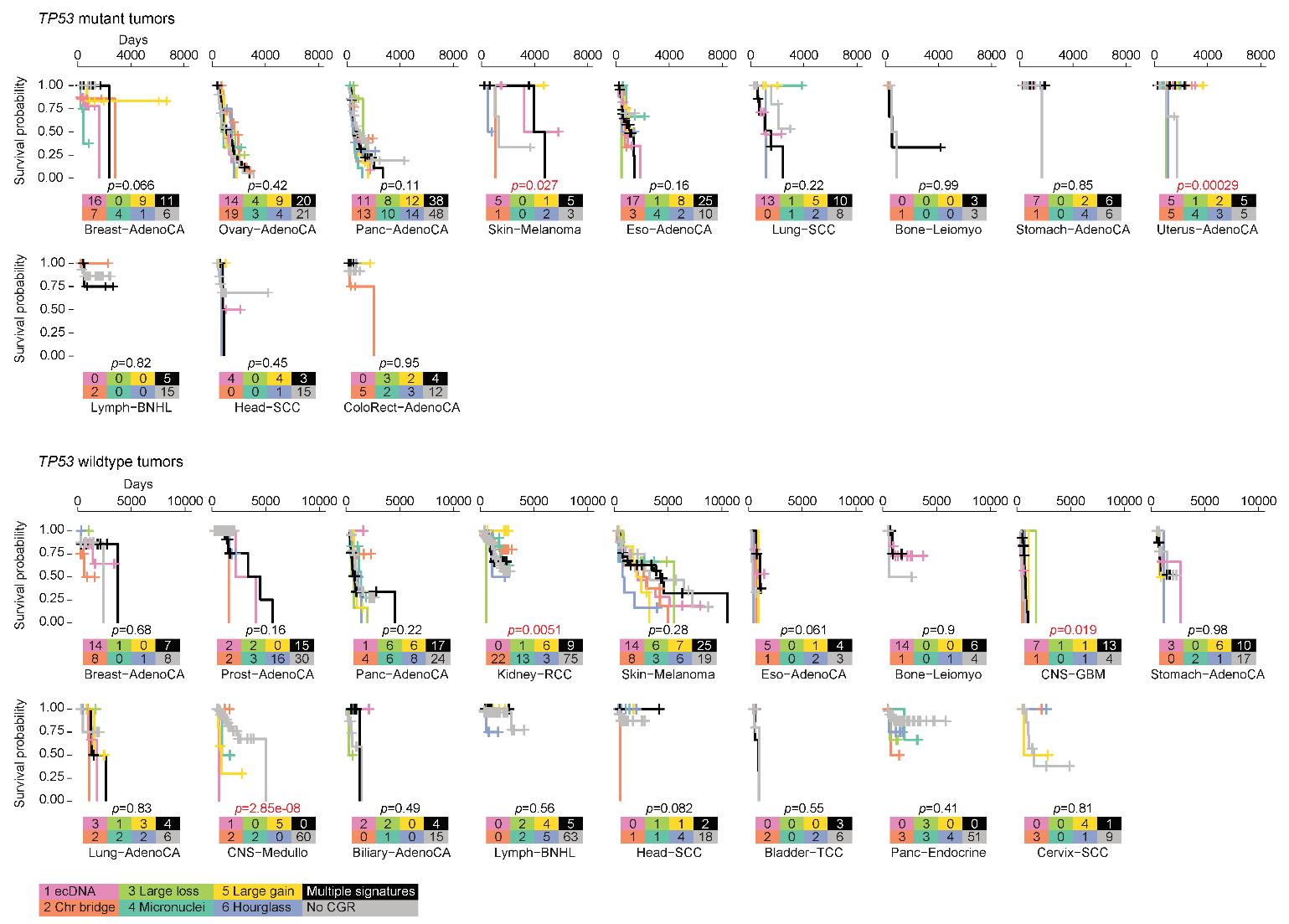

**Figure S8. Comparisons of CGR signatures in patient survival stratified by *TP53* mutation status and tumor type.** Only tumor types with at least three samples in no CGR group and at least three samples in any one of the CGR signature group are displayed. Although *p* values in several tumor types are less than 0.05 (marked in red), they are mainly driven by small sample size with extreme values. In most tumor types, there are no significant differences in survival for patients with or without CGRs in their tumors.

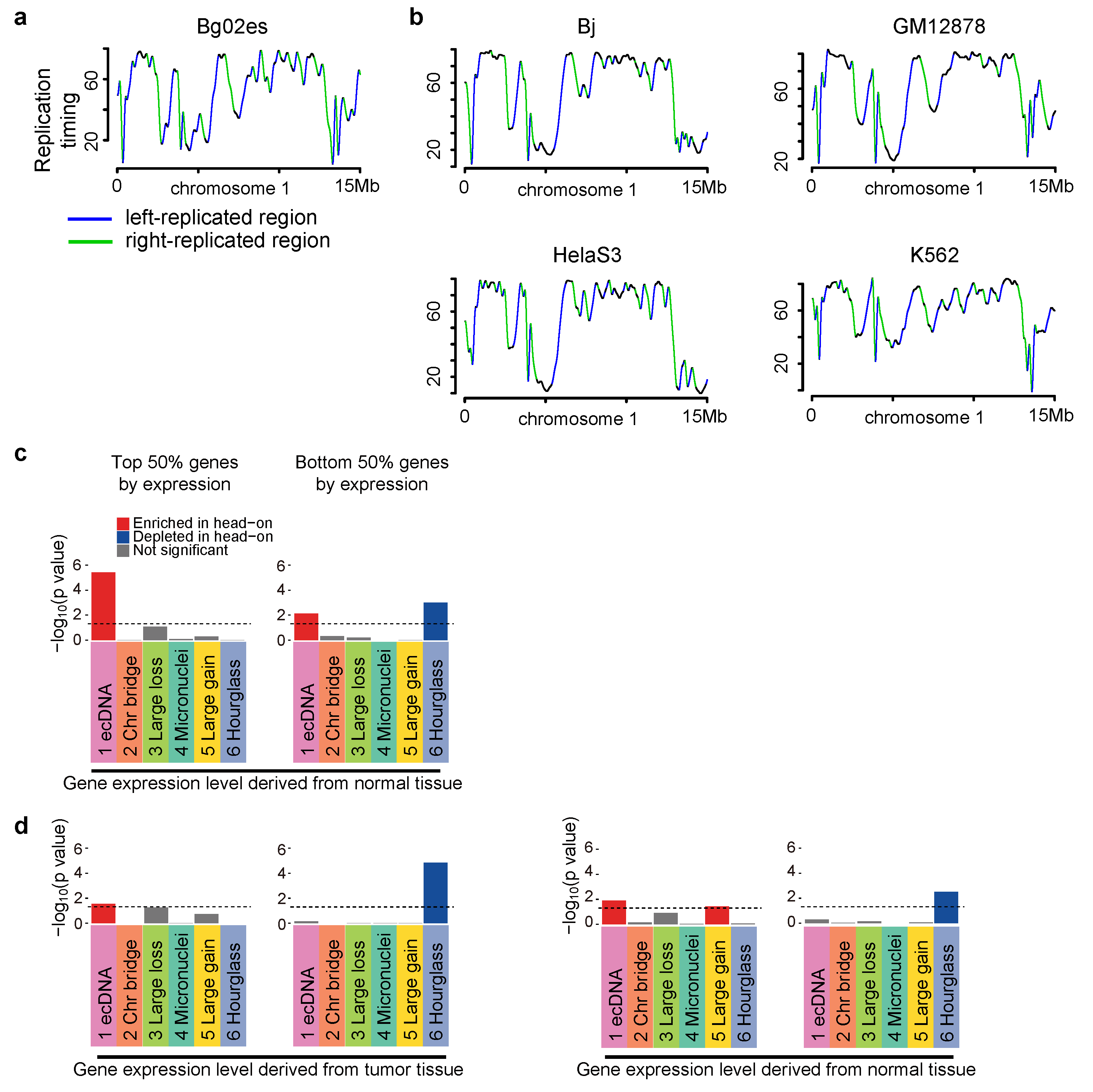

**Figure S9. Transcription-replication collision and CGR breakpoints. a,** Defining DNA replication orientations based on replication-timing profile of Bg02es cell line. **b,** Replication timing profiles and replication orientations in four other cell lines (Bj, GM12878, HelaS3 and K562). **c,** Breakpoint biases in six CGR signatures stratified by gene expression level using gene expression values from normal tissues. **d,** Breakpoint biases in six signatures stratified by gene expression level using gene expression values from tumor and normal tissues. CGR breakpoints within 3 Mb of hotspots are excluded. In **c** and **d**, *p* values are calculated by comparing observed breakpoints and randomly shuffled breakpoints in head-on and co-directional collision regions using Chi-square tests and Bonferroni corrections are performed. Dashed lines represent 0.05 *p* value cutoff.

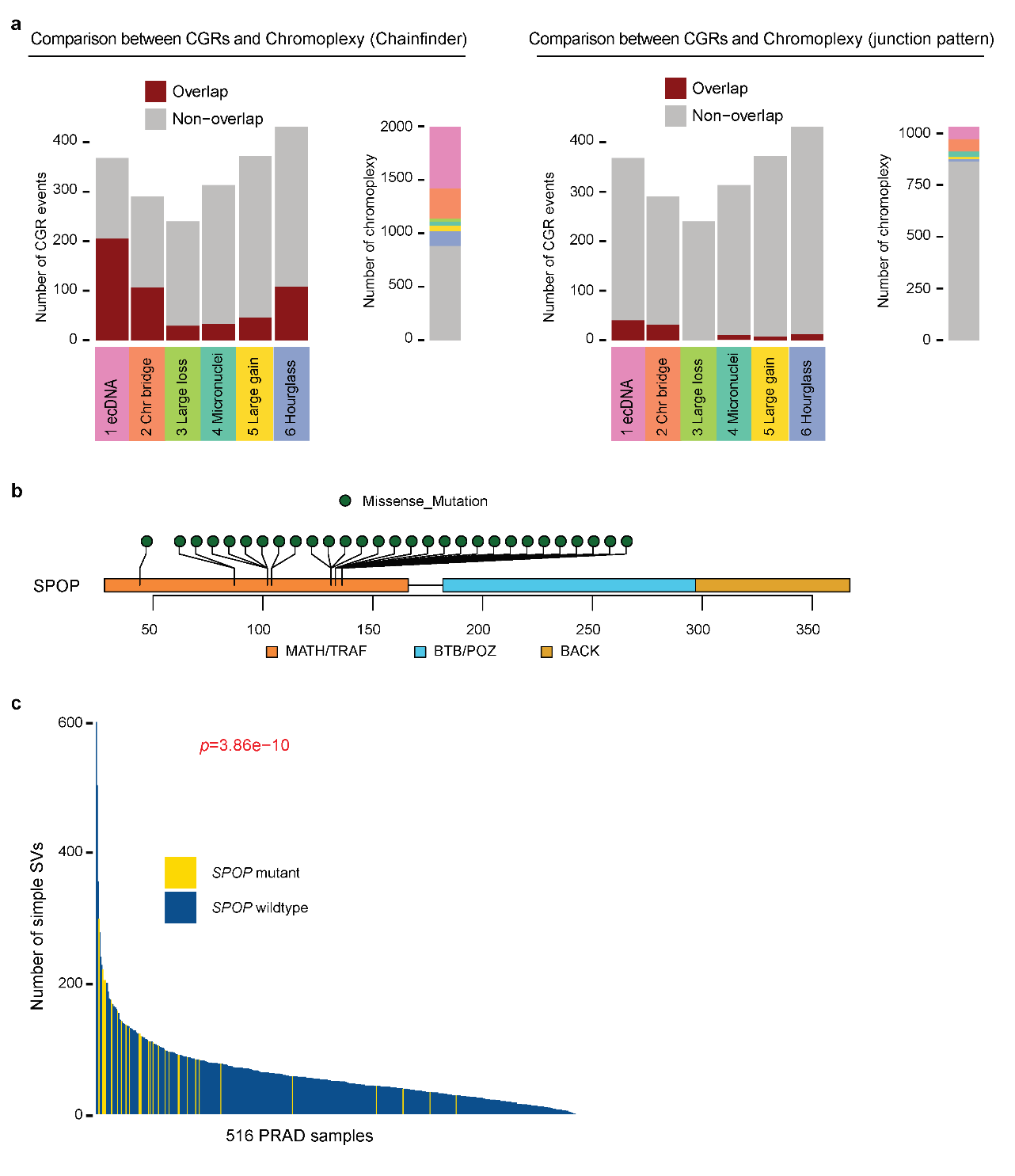

**Figure S10. Comparisons between hourglass chromothripsis and chromoplexy as well as *SPOP* mutations in prostate cancer. a,** Six CGR signatures compared to ChainFinder-predicted and junction-pattern-predicted chromoplexy events. **b,** Somatic mutation distribution in *SPOP* gene in prostate cancer. All mutations are missense mutations in the MATH/TRAF domain which is the target binding domain. **c,** Number of simple SVs in prostate cancers with and without *SPOP* mutations. *P* value is calculated by Wilcoxon rank sum test.

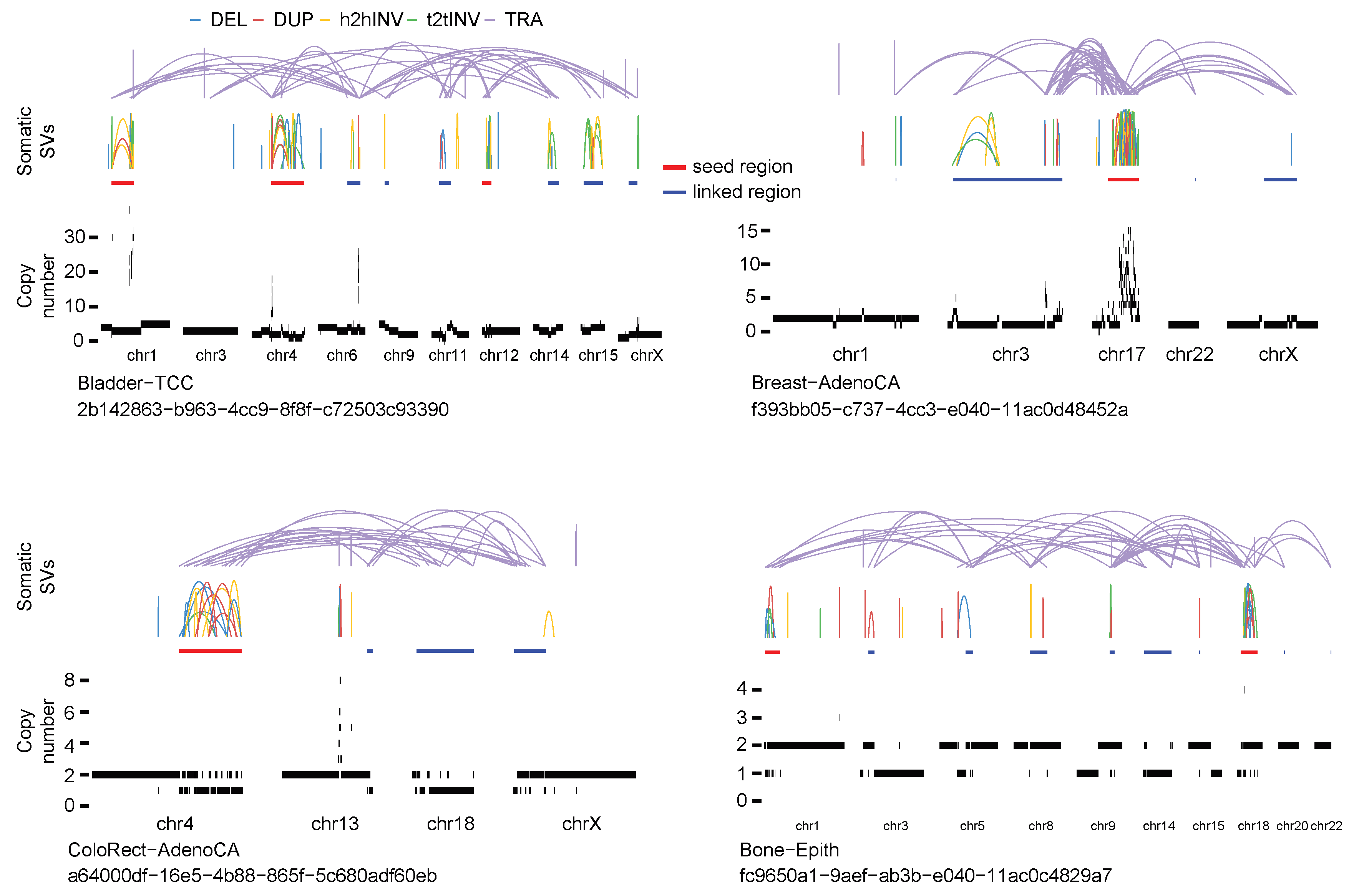

**Figure S11. Identification of CGR seed and linked regions.**
